# mtDNA replication in dysfunctional mitochondria promotes deleterious heteroplasmy via the UPR^mt^

**DOI:** 10.1101/2020.09.01.274472

**Authors:** Qiyuan Yang, Pengpeng Liu, Nadine S. Anderson, Tomer Shpilka, YunGuang Du, Nandhitha Uma Naresh, Kevin Luk, Josh Lavelle, Rilee D. Zeinert, Peter Chien, Scot A. Wolfe, Cole M. Haynes

## Abstract

The accumulation of deleterious mitochondrial genomes (ΔmtDNAs) underlies inherited mitochondrial diseases and contributes to the aging-associated decline in mitochondrial function. In heteroplasmic cells, oxidative phosphorylation (OXPHOS) function declines as the population of ΔmtDNAs increase relative to wildtype mtDNAs. In response to mitochondrial perturbations, the bZIP protein ATFS-1 induces a transcription program to promote the recovery of mitochondrial function. Paradoxically, ATFS-1 is also required to maintain ΔmtDNAs in heteroplasmic worms. However, the mechanism(s) by which ATFS-1 promotes ΔmtDNA accumulation relative to wildtype mtDNAs is unclear. Here, we show that mitochondrial-localized ATFS-1 binds almost exclusively to ΔmtDNAs in heteroplasmic worms. Moreover, we demonstrate that mitochondrial ATFS-1 promotes the preferential binding of the mtDNA replicative polymerase (POLG) to ΔmtDNAs. Interestingly, inhibition of the mtDNA-bound protease LONP-1 increased ATFS-1 and POLG binding to wildtype mtDNAs. Furthermore, LONP-1 inhibition in *C. elegans* and human cybrid cells improved the heteroplasmy ratio and restored OXPHOS function. Our findings suggest that ATFS-1 promotes mtDNA replication by recruiting POLG to mtDNA in a manner that is antagonized by LONP-1. We speculate that this mechanism promotes the repair and expansion of the mitochondrial network by synchronizing mtDNA replication with UPR^mt^ activation driven by nuclear ATFS-1 activity. However, this repair mechanism cannot resolve OXPHOS defects in mitochondria harboring ΔmtDNAs, resulting in an accumulation of ATFS-1 in dysfunctional mitochondria and constitutive replication of ΔmtDNAs.

## INTRODUCTION

Mitochondria provide numerous metabolic functions including being the site of energy production via oxidative phosphorylation (OXPHOS). Most of the ~1200 proteins comprising the mitochondrial proteome are encoded by nuclear genes and must be imported into each mitochondrion following synthesis on cytosolic ribosomes^1^. However, 13 essential OXPHOS components alongside the tRNAs and rRNAs required for their synthesis are encoded by mitochondrial genomes (mtDNAs), which reside in the mitochondrial matrix. Each mitochondrion harbors at least one mtDNA, and most cells harbor 100s-1,000s of mtDNAs.

Mitochondrial dysfunction accumulates as cells age and is accelerated in multiple age-associated diseases, including Alzheimer’s Diseases and Parkinson’s Disease. However, a variety of mitochondrial syndromes or diseases are caused by inherited mutations that impair OXPHOS function. The disease causing mutations can occur in genes required for OXPHOS encoded by either the nuclear genome, or mtDNAs, which acquire mutations at a significantly higher rate than the nuclear genome^2^. A number of single nucleotide variants and deletions are associated with inherited mitochondrial diseases and syndromes affecting ~1:4000 individuals^3^. Because of the high number of mtDNAs per cell, a single mutant mtDNA has little impact. To cause the OXPHOS dysfunction that underlies mitochondrial diseases, the mutant mtDNA must accumulate to ~60% of the total cellular mtDNAs. The mixture of mutant mtDNAs and wildtype mtDNAs is known as heteroplasmy. Several studies using mitochondrial-targeted nucleases that specifically cleave mutant mtDNAs have indicated that a relatively modest reduction in the percentage of ΔmtDNAs is sufficient to improve mitochondrial function in heteroplasmic cells and mouse tissues^4–6^.

The initial mtDNA mutation or deletion likely occurs because an error in mtDNA replication^7^. Two mechanisms are thought to contribute to the “clonal expansion” of the ΔmtDNA. In dividing cells, random, or non-selective, genetic drift can disproportionately increase either genome. An alternative model suggests that large mtDNA deletions allow for quicker replication simply because these genomes are smaller and thus replicate more rapidly^7, 8^. Consistent with these models, mutations that reduce the function of the replicative mtDNA polymerase POLG cause a preferential depletion of mutant mtDNAs and improve heteroplasmy^6,9^. However, the underlying mechanism(s) that drive the clonal amplification of ΔmtDNAs and maintain them at a high enough percentage to cause OXPHOS defects in mitochondrial diseases^10–13^, aging^14^ and Parkinson’s Disease^15^ remain unresolved.

Previously, the bZIP protein ATFS-1 was shown to be required to maintain deleterious heteroplasmy in *C. elegans*^9, 16^. ATFS-1 harbors both a mitochondrial targeting sequence (MTS) and a nuclear localization sequence (NLS) (Fig. 1a) and regulates a transcriptional program known as the mitochondrial unfolded protein response (UPR^mt^)^17^. Under basal conditions, the majority of ATFS-1 is imported into mitochondria, where it is degraded by the protease LONP-1. Mitochondrial dysfunction reduces mitochondrial import capacity, resulting in a percentage of ATFS-1 trafficking to the nucleus, where it activates a transcriptional program that re-establishes mitochondrial homeostasis and promotes mitochondrial biogenesis^18^. Importantly, both nuclear and mitochondrial accumulation of ATFS-1 are required for development during mitochondrial dysfunction^9, 19^. However, the function of ATFS-1 within mitochondria is unclear.

**Fig. 1.**
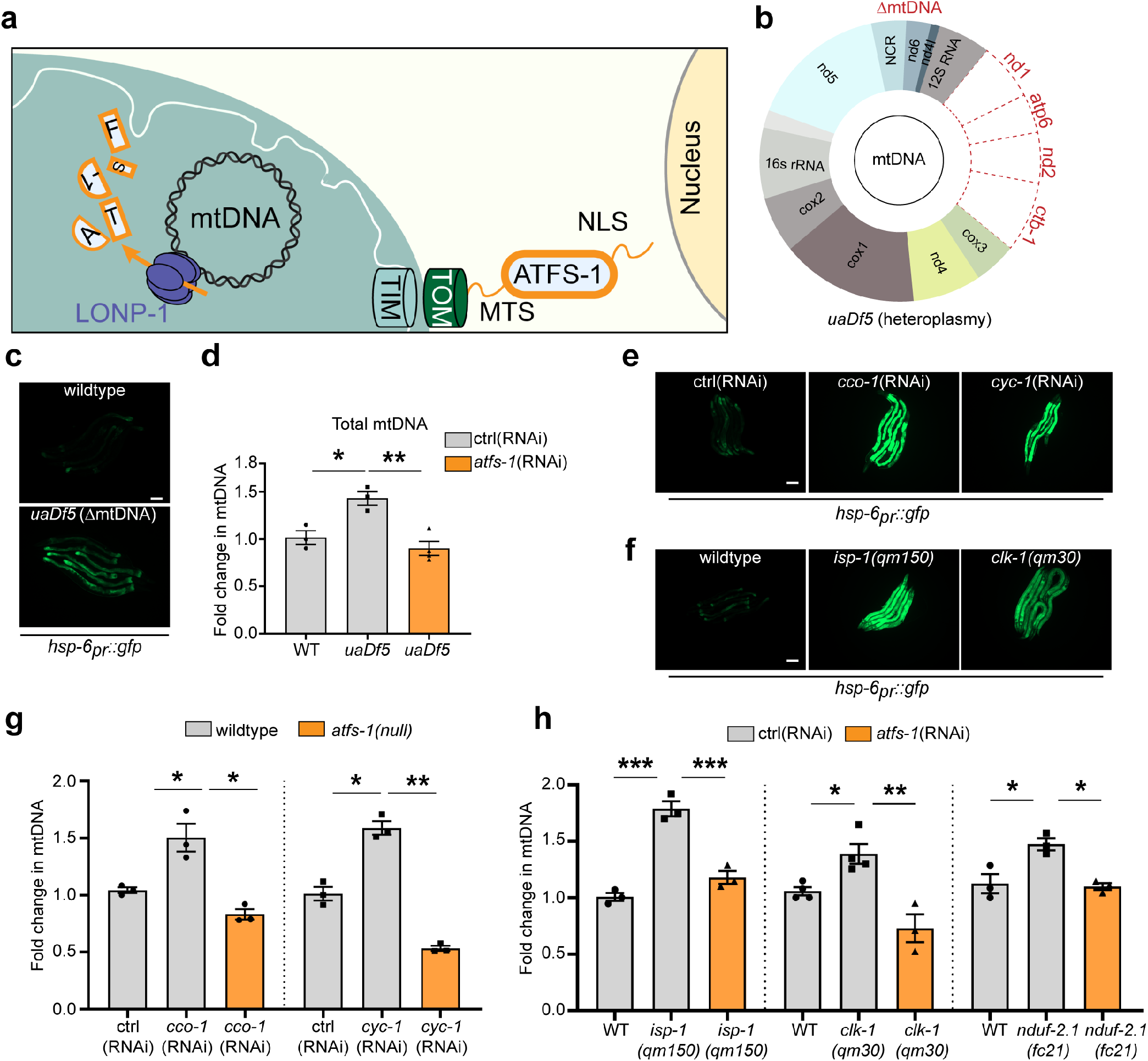
OXPHOS dysfunction increases mtDNAs through ATFS-1. **a**, ATFS-1/UPR^mt^ signaling schematic in healthy cells. **b**, Schematic comparing *C.elegans* wildtype homoplasmic and ΔmtDNA (*uaDf5* deletion) strains. **c**, Photomicrographs of *hsp-6_pr_::gfp* and *hsp-6_pr_::gfp;uaDf5* worms (Scale bar 0.1 mm). **d**, Quantification of total mtDNA in homoplasmic wildtype, *uaDf5* worms and *uaDf5* worms raised on *atfs-1*(RNAi) (*n* = 3). **e**, Photomicrographs of *hsp-6_pr_::gfp* worms raised on control (ctrl), *cco-1* or *cyc-1*(RNAi). **f,** photomicrographs of wildtype, *isp-1(qm150)* and *clk-1(qm30);hsp-6_pr_::gfp* worms. (Scale bar 0.1 mm). **g,** Quantification of total mtDNA in homoplasmic wildtype and *atfs-1(null)* worms raised on control(RNAi) or nuclear-encoded OXPHOS genes (*n* = 3). **h**, Quantification of mtDNA in wildtype, *isp-1(qm150)*, *clk-1(qm30)* and *nduf-2.1(fc21)* mutant worms raised on control(RNAi) or *atfs-1*(RNAi) (*n* = 3-4). (“*n*” means samples from independent biological replicates and each sample contains 40-60 animals; every dot stands for averaged value from 3 technical replicates); One-way ANOVA was used in **d**, **g** and **h**; error bars mean ± S.E.M. *p<0.05, **p<0.01, ***p<0.001.

Here, we report that the maintenance of deleterious heteroplasmy requires the accumulation of ATFS-1 within mitochondria. We demonstrate that in a heteroplasmic *C. elegans* strain, ATFS-1 binds predominantly to ΔmtDNAs. Moreover, the replicative polymerase POLG also binds predominantly to ΔmtDNAs, and that this interaction depends on ATFS-1. Lastly, we demonstrate that the mitochondrial protease LONP-1, which degrades ATFS-1 in functional mitochondria^17, 19^, is required to establish the preferential interaction between ATFS-1 and ΔmtDNAs in heteroplasmic worms. Our findings in *C. elegans* are conserved in cultured human cells as we found that inhibition of LONP1 by siRNA or the drug CDDO^20^, improves heteroplasmy and rescues OXPHOS function in heteroplasmic cells harboring either a mutant mtDNA with a large deletion or with a mutant mtDNA harboring a single base pair substitution.

## RESULTS

### OXPHOS perturbations increase mtDNA quantity

OXPHOS proteins are encoded by genes located within the mitochondria and the nucleus. The heteroplasmic *C. elegans* strain *uaDf5* harbors approximately 40% wildtype mtDNAs and 60% ΔmtDNAs that lack four essential OXPHOS protein coding genes (Fig. 1b)^21^. As shown previously, *uaDf5* worms have impaired respiration^22^, constitutive UPR^mt^ activation, as determined by increased expression of the *hsp-6_pr_::gfp* UPR^mt^ reporter^22, 23^ (Fig. 1c), and an increase in total mtDNAs (Fig. 1d)^21^.

To examine the impact of OXPHOS perturbation on mtDNA content, we impaired several OXPHOS components in wildtype homoplasmic worms. As expected, worms raised on *cco-1*(RNAi) (complex IV) or *cyc-1*(RNAi) (cytochrome c) had increased *hsp-6_pr_::gfp* activation, as did strains with loss of function mutations *isp-1(qm150)* (complex III) or *clk-1(qm30)* (ubiquinone biosynthesis) (Figs. 1e,f). Intriguingly, each of the OXPHOS perturbations resulted in increased mtDNA content as determine by qPCR (Figs. 1g,h and Supplementary Fig. 1a), which is consistent with previous reports^24, 25^. Importantly, the increase in mtDNAs caused by OXPHOS perturbation was impaired in *atfs-1(null)* worms that lacks the entire *atfs-1* open reading frame^26^ (Fig. 1g) as well as worms raised on *atfs-1*(RNAi) (Fig. 1h). Similarly, the increase in total mtDNAs in heteroplasmic worms was also reduced when raised on *atfs-1*(RNAi) (Fig. 1d). Combined, these findings indicate that the increased mtDNA content in both homoplasmic and heteroplasmic worms caused by OXPHOS perturbation requires *atfs-1*.

### Increased POLG-mtDNA binding during OXPHOS perturbation requires ATFS-1

We next sought to gain insight into the mechanism by which mtDNAs are increased during OXPHOS perturbation. We previously found that ATFS-1 accumulates within mitochondria when mitochondrial function is perturbed by inhibiting the essential mitochondrial protease SPG-7 or by raising worms in the presence of ethidium bromide, which impairs mtDNA replication^17, 27^. Intriguingly, ChIP-sequencing indicated that during mitochondrial dysfunction, ATFS-1 binds mtDNAs at a single site within the non-coding region (NCR)^28^ (Fig. 2a), which in mammals contains sequence elements that regulate mtDNA replication^29^. Here, we found that ATFS-1 also accumulates in the mitochondrial fraction of worms raised on *cco-1*(RNAi) (Fig. 2b). Furthermore, ATFS-1 ChIP followed by qPCR indicated that ATFS-1 also interacts with mtDNA when worms are raised on *cco-1*(RNAi), which was impaired in *atfs-1(null)* worms (Fig. 2c). Combined these data suggest that ATFS-1 accumulates within mitochondria and interacts with mtDNAs when OXPHOS is impaired.

**Fig. 2.**
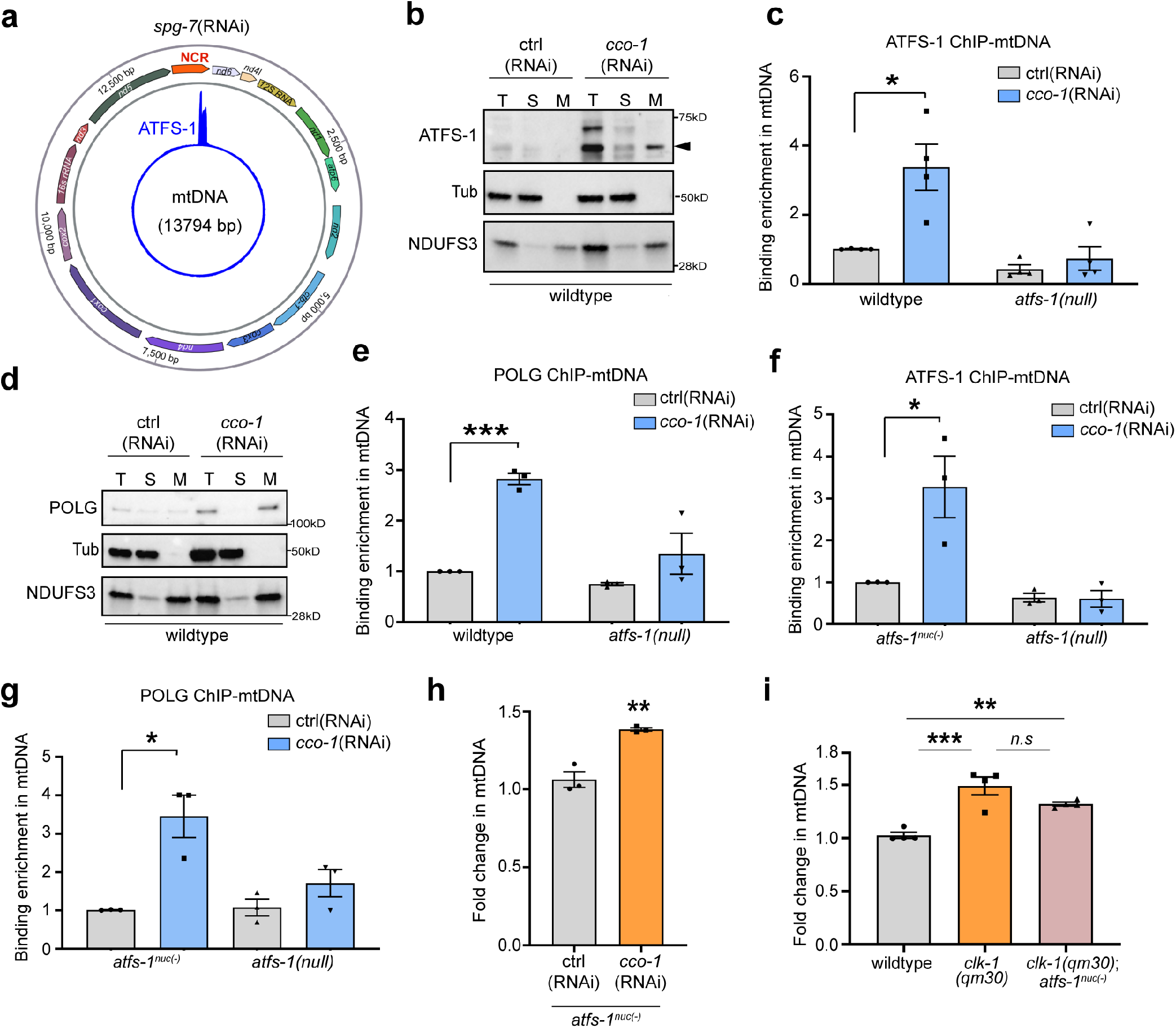
Defective OXPHOS impedes the degradation ATFS-1 and facilitates mitochondrial localized ATFS-1 and POLG binding to mtDNAs. **a**, ATFS-1 ChIP-seq profile of the entire mitochondrial genome (mtDNA) of wildtype worms raised on *spg-7*(RNAi) (adapted from^19^). **b**, ATFS-1 immunoblots of wildtype worms raised on control or *cco-1*(RNAi) after fractionation into total lysate (T), post-mitochondrial supernatant (S), and mitochondrial pellet (M). Tubulin (Tub) and the OXPHOS component (NDUFS3) are used as loading controls. Arrow is mitochondrial-localized ATFS-1. **c**, Quantification of mtDNA following ATFS-1 ChIP-mtDNA in homoplasmic wildtype worms and homoplasmic *atfs-1(null)* worms raised on control or *cco-1*(RNAi) (*n* = 4). **d**, POLG immunoblots of wildtype worms raised on control or *cco-1*(RNAi) after mitochondrial fractionation. **e**, Quantification of total mtDNA following POLG ChIP-mtDNA in wildtype or *atfs-1(null)* homoplasmic worms (*n* = 3). **f**, Quantification of mtDNA following ATFS-1 ChIP-mtDNA in homoplasmic *atfs-1^nuc(−)^* worms and homoplasmic *atfs-1(null)* worms raised on control or *cco-1*(RNAi) (*n* = 3). **g**, Quantification of total mtDNA following POLG ChIP-mtDNA in homoplasmic *atfs-1^nuc(−)^* and *atfs-1(null)* worms raised on control or *cco-1*(RNAi) (*n* = 3). **h**, Quantification of total mtDNA in *atfs-1^nuc(−)^* homoplasmic worms raised on control(RNAi) or *cco-1*(RNAi)) (*n* = 3). **i**, Quantification of total mtDNA in wildtype, *clk-1(qm30)* and *clk-1(qm30)*;*atfs-1^nuc(−)^* homoplasmic wildtype worms (*n* = 4, One-way ANOVA). (“*n*” means independent biological replicates and each sample pooled from large population in **c**, **e-g**; contains 40-60 animals in **h** and **i**; every dot stands for averaged value from 3 technical replicates); Two-tailed Student’s t test was used except to **i**; error bars mean ± S.E.M. *p<0.05, **p<0.01, ***p<0.001.

To further explore the relationship between ATFS-1 and mtDNA accumulation during OXPHOS perturbation, we generated POLG antibodies which detected a ~120 KD band that co-fractionated with the OXPHOS protein NDUFS3 (Supplementary Fig. 1b) and was depleted by *polg*(RNAi) (Supplementary Fig. 1c). Similar to ATFS-1, POLG protein levels increased within mitochondria when raised on *cco-1*(RNAi) (Fig. 2d), and POLG interacted with more mtDNAs (Fig. 2e). To determine if the increased POLG-mtDNA interaction during OXPHOS perturbation required *atfs-1*, we also performed POLG ChIP-mtDNA in wildtype and *atfs-1(null)* worms raised on *cco-1*(RNAi). Interestingly, the increased POLG-mtDNA interaction in wildtype worms raised on *cco*-1(RNAi) was impaired in *atfs-1(null)* worms, suggesting that ATFS-1 is required for increased POLG-mtDNA binding during OXPHOS dysfunction (Fig. 2e).

Our data indicate that ATFS-1 is required for POLG to bind mtDNA during OXPHOS perturbation. We next sought to examine if the increased POLG-mtDNA binding required nuclear-localized ATFS-1. To circumvent the potentially confounding effects of the induction of *polg* mRNA by ATFS-1 during mitochondrial stress^9, 17^, we first generated the *atfs-1^nuc(−)^* strain in which the NLS within ATFS-1 was impaired via genome editing (Supplementary Figs. 2a,b). Importantly, the *atfs-1^nuc(−)^* strain was unable to induce *hsp-6* mRNA when raised on *spg-7*(RNAi), consistent with impaired nuclear function (Supplementary Fig. 2c). Furthermore, we introduced the *atfs-1^nuc(−)^* mutation into *atfs-1(et18)* worms, which causes constitutive induction of the UPR^mt^ reporter *hsp-6_pr_::gfp* and *hsp-6* mRNA due to a mutation that perturbs the MTS of ATFS-1^30^. The *atfs-1^nuc(−)^* mutation in *atfs-1(et18)* worms suppressed activation of the UPR^mt^ reporter *hsp-6_pr_::gfp*, and *hsp-6* mRNA (Supplementary Figs. 2e-g). Induction of *hsp-6* and *polg* mRNA when raised on *cco-1*(RNAi) was also impaired in *atfs-1^nuc(−)^* worms consistent with impaired nuclear transcription (Supplementary Figs. 2g,h). Furthermore, unlike in wildtype worms, POLG protein levels were not increased in *atfs-1^nuc(−)^* worms indicating that *atfs-1^nuc(−)^* worms were unable to regulate nuclear transcription during OXPHOS perturbation (Supplementary Figs. 2h,i). Importantly, ATFS-1^nuc(−)^ protein accumulated within mitochondria to similar levels as wildtype ATFS-1 upon LONP-1 inhibition (Supplementary Fig. 2j) indicating the protein was expressed and processed similarly to wildtype ATFS-1. Lastly, ATFS-1^nuc(−)^ bound similar amounts of mtDNA as wildtype ATFS-1 when raised on *cco-1*(RNAi) as determined by ChIP (Figs. 2c,f).

To determine if the nuclear function of ATFS-1 is required for POLG to bind mtDNAs during OXPHOS perturbation, we examined the POLG-mtDNA interaction in *atfs-1(null)* and *atfs-1^nuc(−)^* worms raised on control or *cco-1*(RNAi). Strikingly, significantly more POLG was bound to mtDNA in *atfs-1^nuc(−)^* worms relative to *atfs-1(null)* worms (Fig. 2g), suggesting that the *atfs-1-*dependent increase in POLG-mtDNA binding during OXPHOS perturbation does not require the nuclear activity of ATFS-1. mtDNA content also increased in *atfs-1^nuc(−)^* worms upon OXPHOS perturbation caused by *cco-1*(RNAi) (Fig. 2h) as well as in *clk-1(qm30)* worms (Fig. 2i), similar to ATFS-1 wildtype worms. Combined, these results suggest that the accumulation of ATFS-1 within mitochondria during OXPHOS perturbation is sufficient to increase POLG-mtDNA binding and mtDNA content.

### Degradation of ATFS-1 by LONP-1 within mitochondria impairs mtDNA propagation

We next sought to determine the events leading to the accumulation of ATFS-1 within mitochondria when OXPHOS is perturbed. During normal growth, the ATP-dependent protease LONP-1 degrades the majority of ATFS-1 once the bZIP protein has been imported into the mitochondrial matrix^17^. LONP-1 is an ATP-dependent protease known to recognize and degrade mitochondrial proteins damaged by reactive oxygen species^31^. LONP-1 has also been shown to interact with mtDNA in diverse species (Fig. 1a)^32, 33^, and regulate mtDNA replication^34, 35^.

To further examine the interaction between LONP-1 and mtDNA in *C. elegans*, we generated a strain in which the C-terminus of LONP-1 was tagged with the FLAG epitope via genome editing (Fig. 3a and Supplementary Fig. 3a). Introduction of the FLAG epitope did not impair worm development or cause UPR^mt^ activation, suggesting it did not adversely affect LONP-1 function (Supplementary Figs. 3b,c). Consistent with findings in other species^32, 33^, LONP-1 interacted with mtDNA in *C. elegans* as determined by LONP-1^FLAG^ ChIP-mtDNA qPCR (Fig. 3b). We next examined where in mtDNA LONP-1^FLAG^ binds in wildtype homoplasmic worms. LONP-1^FLAG^ ChIP-seq indicated that the protease was enriched at several G-rich sites throughout mtDNA (Fig. 3c), but was especially enriched within the NCR (Fig. 3d). Interestingly, the strongest LONP-1^FLAG^ peak within the NCR overlapped with the ATFS-1 binding site (Fig. 3d and Supplementary Fig. 3d) suggesting a potential role in mediating the ATFS-1-mtDNA interaction.

**Fig. 3.**
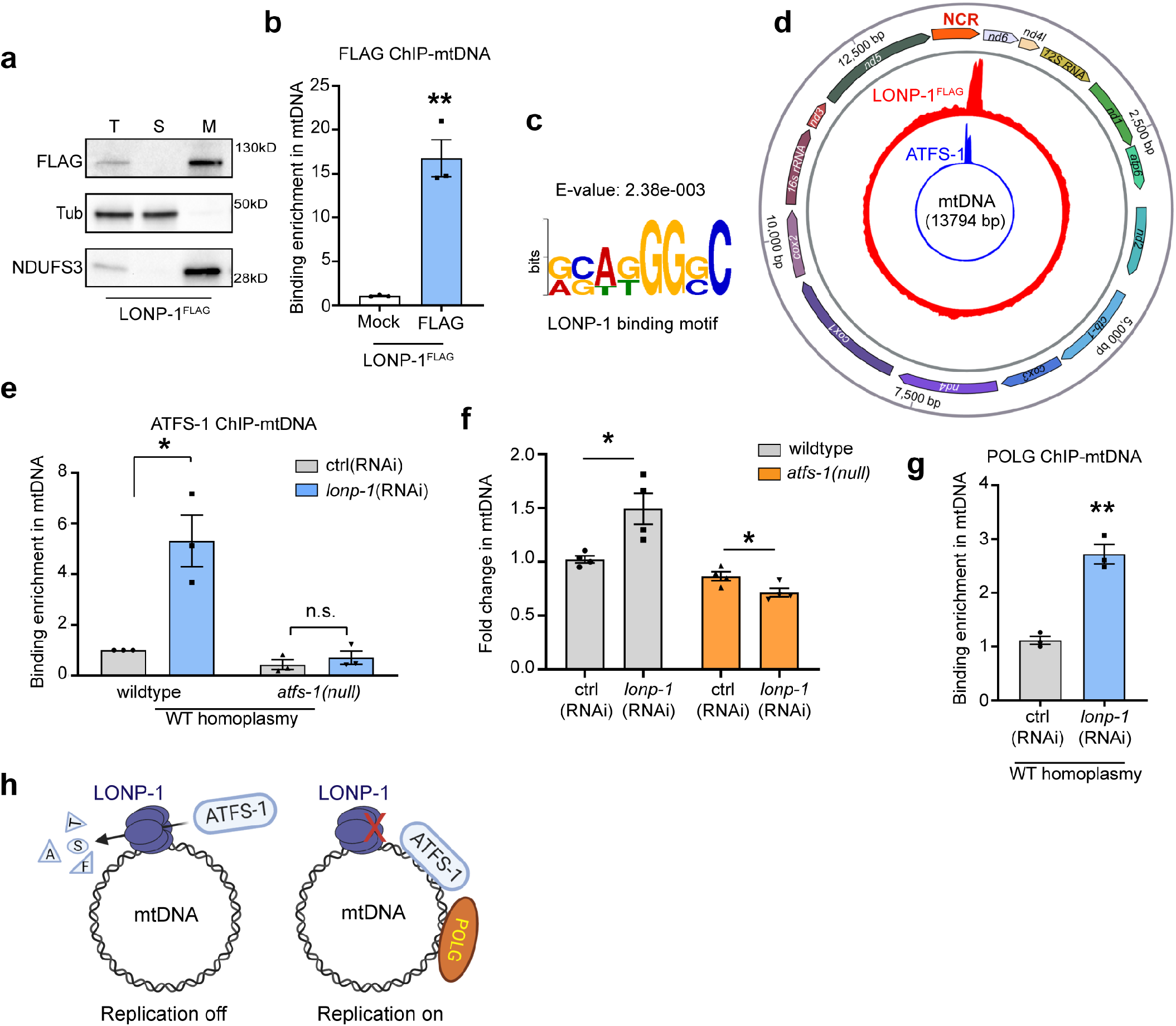
LONP-1 limits ATFS-1 binding to wildtype mtDNAs and impairs replication. **a**, FLAG immunoblots of LONP-1^FLAG^ worms following fractionation into total lysate (T), post-mitochondrial supernatant (S), and mitochondrial pellet (M). Tubulin (Tub) and the OXPHOS component NDUFS3 are loading controls. **b**, Quantification of mtDNA from homoplasmic LONP-1^FLAG^ worms following ChIP-mtDNA using FLAG or control (Mock) antibody (*n* = 3). **c**, LONP-1 consensus binding motif within mtDNA. **d**, ChIP-seq profile of mtDNA from homoplasmic LONP-1^FLAG^ worms raised on control(RNAi) using FLAG antibody (red). ATFS-1 ChIP-seq profile from homoplasmic worms raised on *spg-7*(RNAi) (blue) (as in Fig.2a). **e**, Quantification of total mtDNA following ATFS-1 ChIP-mtDNA in wildtype or *atfs-1(null)* homoplasmic worms raised on control(RNAi) or *lonp-1*(RNAi) (*n* = 3). **f**, Quantification of mtDNA in homoplasmic wildtype and *atfs-1(null)* worms raised on control(RNAi) or *lonp-1*(RNAi) (*n* = 4). **g**, Quantification of total mtDNA following POLG ChIP-mtDNA in wildtype homoplasmic worms raised on control(RNAi) or *lonp-1*(RNAi) (*n* = 3). **h**, Schematic of the relationship between LONP-1 activity, mitochondrial ATFS-1 accumulation and mtDNA replication. (“*n*” means independent biological replicates and each sample pooled from large populations in **b**, **e**, **g**; each sample contains 40-60 animals in **f**; every dot stands for averaged value from 3 technical replicates); Two-tailed Student’s t test was used; error bars mean ± S.E.M. *p<0.05, **p<0.01.

We next examined the impact of *lonp-1*(RNAi) on mtDNA accumulation in homoplasmic wildtype worms. When raised on *lonp-1*(RNAi), the binding of ATFS-1 to mtDNA increased by ~6-fold (Fig. 3e) which correlated with an increase in mtDNA content (Fig. 3f). Importantly, exposure to *lonp-*1(RNAi) also increased the binding of POLG to mtDNA (Fig. 3g). Moreover, LONP-1 inhibition increased mtDNA content in *atfs-1^nuc(−)^* worms (Supplementary Fig. 3e), but not in *atfs-1(null)* worms (Fig. 3f). In short, these findings support a role for mitochondrial-localized ATFS-1 in promoting mtDNA replication which is impaired by LONP-1-dependent degradation (Fig. 3h). While it remains to be determined how LONP-1 mediates ATFS-1 accumulation, we propose that 1) the decrease of ATP within perturbed mitochondria or 2) the increased accumulation of misfolded proteins that are the primary substrates of the protease^36^ may impair LONP-1 function and allow ATFS-1 to accumulate within dysfunctional mitochondria.

### Maintenance of deleterious heteroplasmy does not require ATFS-1 nuclear activity

We and the Patel Lab previously found that ATFS-1 is required to maintain ΔmtDNAs^9, 16^ in a heteroplasmic worm strain^21^. Here, we crossed the *atfs-1(null)* allele into *uaDf5* heteroplasmic worms. Consistent with previous results, ΔmtDNAs were severely depleted in heteroplasmic *atfs-1(null)* worms (Fig. 4a). The loss of ΔmtDNAs in the absence of *atfs-1* could either be due to increased mitophagy of mitochondria harboring ΔmtDNAs^16, 37^, or decreased replication of ΔmtDNAs. To examine the role of mitophagy, we generated heteroplasmic strains lacking the mitophagy component Parkin (PDR-1 in worms) (Supplementary Fig. 4a). As expected, *pdr-1*-deficient worms had increased ΔmtDNAs relative to wildtype worms consistent with mitophagy limiting the accumulation of ΔmtDNAs^22, 23^. However, *atfs-1(null);pdr-1(tm598)* worms had a significant reduction in ΔmtDNAs relative to *pdr-1(tm598)* worms, indicating that ATFS-1 promotes heteroplasmy via a mechanism independent of mitophagy.

**Fig. 4.**
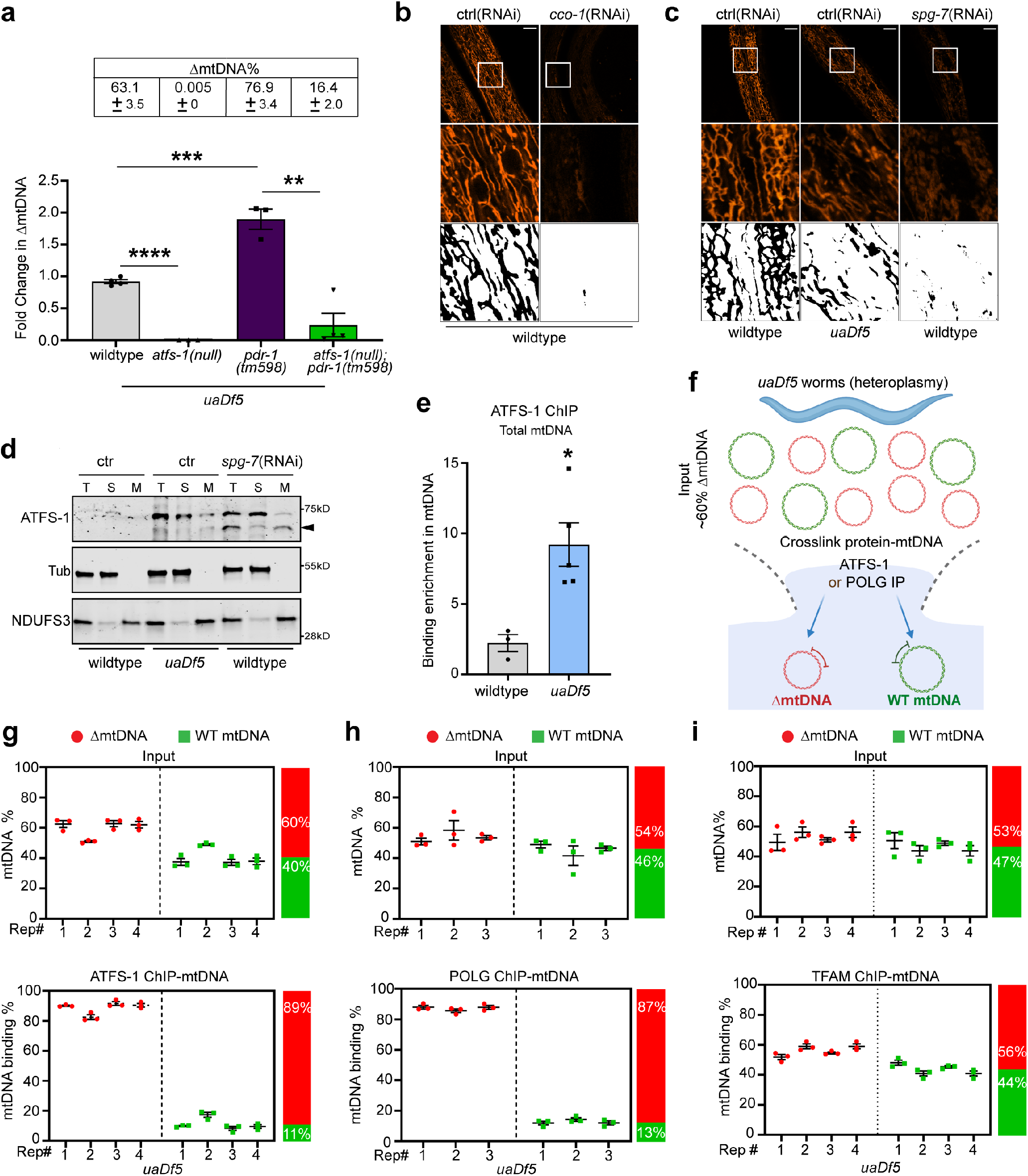
ATFS-1 and POLG primarily interact with ΔmtDNAs in heteroplasmic worms. **a**, ΔmtDNA quantification as determined by qPCR in *uaDf5* worms, *atfs-1(null)*;*uaDf5*, *pdr-1(tm598)*;*uaDf5* and *atfs-1(null);pdr-1(tm598)*;*uaDf5* worms (*n* = 3-4, One-way ANOVA). **b**, Images of TMRE-stained micrographs of wildtype worms raised on control(RNAi) or *cco-1*(RNAi). Scale bar, 10 μM. **c,** Images of TMRE-stained micrographs of heteroplasmic (ΔmtDNA) worms raised on control(RNAi), or wildtype worms raised on control or *spg-7*(RNAi). Scale bar, 10 μM. **d**, Immunoblots of wildtype worms raised on control or *spg-7*(RNAi) and heteroplasmic (ΔmtDNA) worms raised on control(RNAi) after fractionation into total lysate (T), post-mitochondrial supernatant (S), and mitochondrial pellet (M). Tubulin (Tub) and the OXPHOS component (NDUFS3) are used as loading controls. Arrow is mitochondrial-localized ATFS-1. **e**, Quantification of total mtDNA following ATFS-1 ChIP-mtDNA in homoplasmic wildtype or ΔmtDNA worms (*n* = 3 or 5, two-tailed Student’s t test). **f**, Workflow of ATFS-1 or POLG ChIP-mtDNA and quantification of wildtype mtDNA and ΔmtDNA in heteroplasmic worms. **g**, Quantification of wildtype mtDNA and ΔmtDNA by qPCR following ATFS-1 ChIP-mtDNA in heteroplasmic worms. Post-lysis/Input ΔmtDNA ratio was 60% (*n* = 4). **h**, Quantification of wildtype mtDNA and ΔmtDNA by qPCR following POLG ChIP-mtDNA in heteroplasmic worms. Post-lysis/Input ΔmtDNA ratio was 54% (*n* = 3). **i**, Quantification of wildtype mtDNA and ΔmtDNA following TFAM IP-mtDNA in heteroplasmic worms. Post-lysis/Input ΔmtDNA ratio was 53% (*n* = 4). (“*n*” means independent biological repeats and each sample contains 40-60 animals in **a**; each sample pooled from large populations in **e**, **g-i**; every dot stands for averaged value from 3 technical replicates in **a**, **e** or strands for every technical replicates in **g-i**); error bars mean ± S.E.M. *p<0.05, **p<0.01, ****p<0.0001.

As OXPHOS perturbation increased mtDNA content in homoplasmic wildtype worms via ATFS-1, we hypothesized that OXPHOS dysfunction may contribute to the increased mtDNA content in heteroplasmic worms (Fig. 1d). We previously found that heteroplasmic worms consumed less oxygen than wildtype worms^9^. To further evaluate mitochondrial function, we examined mitochondrial membrane potential by staining with tetramethylrhodamine ethyl ester (TMRE). As expected, TMRE staining was decreased in heteroplasmic worms relative to wildtype worms, but stronger than in *cco-1*(RNAi)- or *spg-7*(RNAi)-treated worms consistent with intermediate OXPHOS function (Figs. 4b,c and Supplementary Figs. 4b,c). Importantly, ATFS-1 also accumulated within the mitochondrial fraction of heteroplasmic worms, as in worms raised on *spg-7*(RNAi) or *cco-1*(RNAi) (Fig. 4d, also see Fig. 2b). Combined, these results indicate that ATFS-1 accumulates within dysfunctional mitochondria caused by impairment of either a nuclear-encoded OXPHOS component, a nuclear-encoded mitochondrial protease, or in worms harboring ΔmtDNAs.

We next examined if maintenance of ΔmtDNAs required the nuclear activity of ATFS-1 by crossing the *atfs-1^nuc(−)^* allele into the heteroplasmic background. Consistent with impaired nuclear activity, *hsp-6_pr_::gfp* was not increased in heteroplasmic *atfs-1^nuc(−)^* worms (Supplementary Figs. 4d,e). Impressively, unlike *atfs-1(null)* worms, *atfs-1^nuc(−)^* worms were able to maintain ΔmtDNAs to nearly the same level as wildtype *atfs-1* worms (Supplementary Fig. 4f) indicating the role of ATFS-1 in promoting heteroplasmy is largely independent of the nuclear activity. Consistent with the accumulation of ATFS-1 in dysfunctional mitochondria (Figs. 2b and 4d), ATFS-1 bound more total mtDNAs in heteroplasmic worms than in wildtype homoplasmic worms as determined by ChIP-mtDNA (Fig. 4e). Similarly, POLG also interacted with more total mtDNAs in heteroplasmic worms than in wildtype homoplasmic worms suggesting increased mtDNA replication in heteroplasmic worms (Supplementary Fig. 4g). Combined, these findings suggest a role for mitochondrial-localized ATFS-1 in maintaining deleterious mtDNA heteroplasmy.

### In heteroplasmic worms, ATFS-1 and POLG primarily bind to ΔmtDNAs

To gain insight into the mechanism by which ATFS-1 maintains heteroplasmy, we again examined the ATFS-1-mtDNA interaction. Because wildtype mtDNAs and ΔmtDNAs both harbor the ATFS-1 binding site, we sought to determine if ATFS-1 differentially interacted with each genome. The interaction between ATFS-1 and wildtype mtDNAs or ΔmtDNAs was evaluated via qPCR or 3D digital PCR following ATFS-1 ChIP (Fig. 4f)^22^. As before, qPCR of mtDNA from heteroplasmic whole worm lysate indicated the strain harbored ~60% ΔmtDNAs and ~40% wildtype mtDNAs (Fig 1b), (Supplementary Figs. 5a,b). qPCR following ATFS-1 ChIP indicated that of the mtDNAs that interacted with ATFS-1, 90% were ΔmtDNAs and 10% were wildtype mtDNAs, indicating that ATFS-1 is significantly enriched on ΔmtDNAs (Fig. 4g and Supplementary Figs. 5c,d).

Previously, we found that inhibition of POLG caused depletion of ΔmtDNAs in heteroplasmic worms relative to wildtype mtDNAs^9^, which is similar to findings in a *Drosophila* model of deleterious mtDNA heteroplasmy^38^. Both findings suggest increased replication of ΔmtDNAs in heteroplasmic cells. To further explore the relationship between ATFS-1 and mtDNA replication, we performed POLG ChIP-mtDNA. Interestingly, ChIP-mtDNA using the POLG antibody indicated that the replicative polymerase also interacted with ~90% ΔmtDNAs and 10% wildtype mtDNAs (Fig. 4h and Supplementary Figs. 5e,f), similarly to ATFS-1 (Fig. 4g). As a control, we generated antibodies to the mtDNA packaging protein HMG-5 (TFAM in mammals) (Supplementary Figs. 5g,h), which interacts with mtDNAs independent of replication^39^. In contrast to ATFS-1 and POLG, the percentage of ΔmtDNAs bound to HMG-5 reflected the percentage within the whole worm lysate (Fig. 4i). Combined, these data indicate that ATFS-1 and a component of the replisome are enriched on ΔmtDNAs relative to wildtype mtDNAs in heteroplasmic worms, consistent with the mutant mtDNA having a replicative advantage.

### The protease LONP-1 is required to maintain heteroplasmy

Because LONP-1 binds mtDNA and its proteolytic activity prevents ATFS-1 accumulation in functional mitochondria^17^ (See Figs. 3d,e), we examined the role of LONP-1 in heteroplasmy maintenance. We generated antibodies to *C. elegans* LONP-1 that recognized a ~130 KD band that was reduced when raised on *lonp-1*(RNAi) (Supplementary Fig. 6a). Via ChIP-qPCR, we found that LONP-1 binds ~60 fold more mtDNAs than ATFS-1 in wildtype worms (Supplementary Fig. 6b). In heteroplasmic worms, LONP-1 bound similar percentages of wildtype and ΔmtDNAs suggesting that unlike ATFS-1 and POLG-1, LONP-1 interacts with mtDNAs independent of mitochondrial dysfunction (Fig. 5a and Supplementary Fig. 6c). Combined, these data suggest that LONP-1 is constitutively bound to mtDNAs and heteroplasmy is not maintained by uneven mtDNA binding by the protease.

**Fig. 5.**
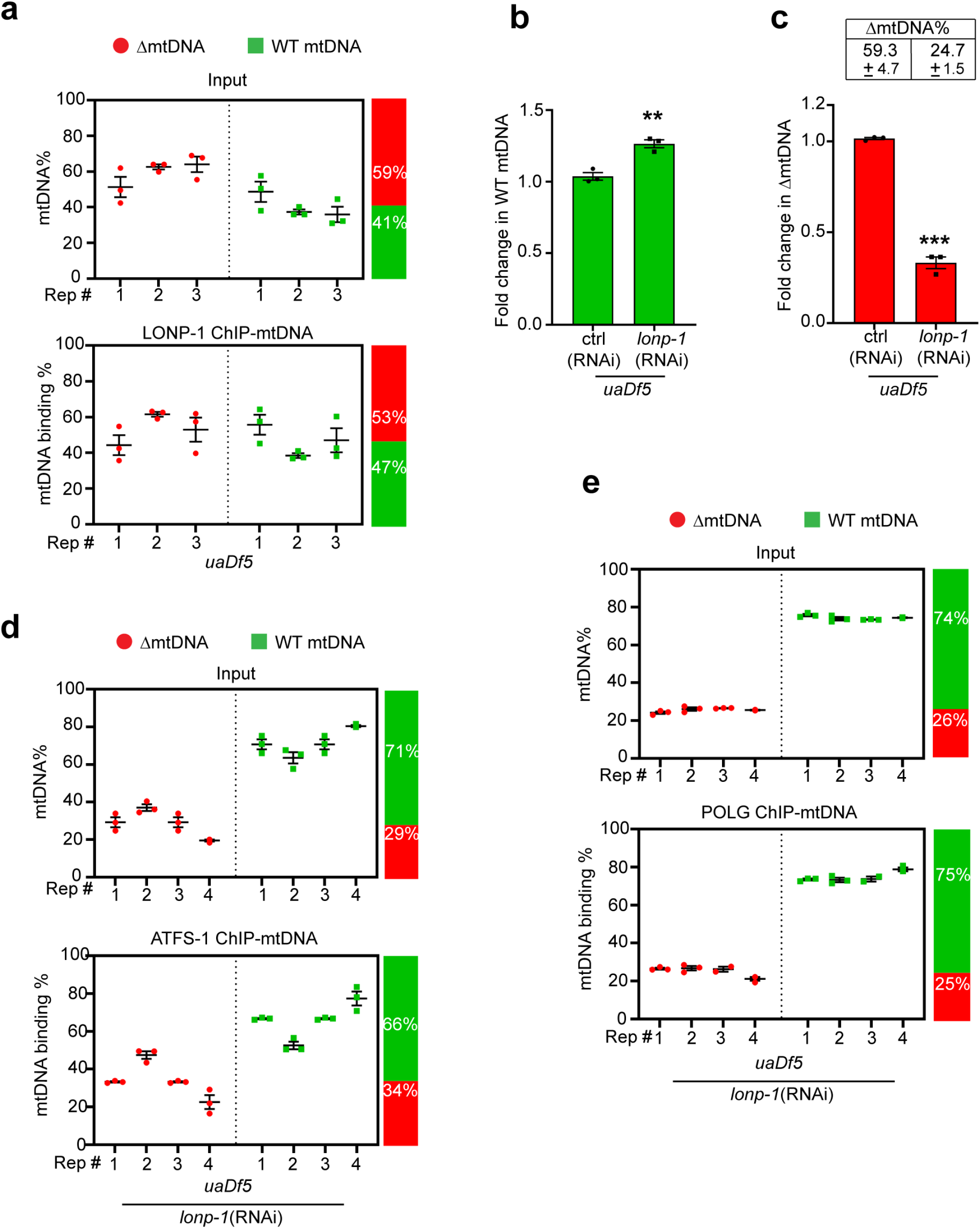
LONP-1 is required to maintain heteroplasmy. **a**, ΔmtDNA and wildtype mtDNA quantification by qPCR following LONP-1 ChIP-mtDNA in heteroplasmic worms. Post-lysis/Input ΔmtDNA ratio was 59% (*n* = 3). **b**, Quantification of wild type mtDNA by qPCR in heteroplasmic worms raised on control(RNAi) or *lonp-1*(RNAi) (*n* = 3). **c**, Quantification of ΔmtDNA by qPCR in heteroplasmic *uaDf5* worms raised on control(RNAi) or *lonp-1*(RNAi) (*n* = 3). **d**, ΔmtDNA and wild-type mtDNA quantification by qPCR following ATFS-1 ChIP-mtDNA in heteroplasmic worms raised on control(RNAi) or *lonp-1*(RNAi) (*n* = 4). **e**, ΔmtDNA and wild-type mtDNA quantification by qPCR following POLG ChIP-mtDNA in heteroplasmic worms raised on *lonp-1*(RNAi). Post-lysis/Input ΔmtDNA ratio was 26% (*n* = 4). (“*n*” means independent biological repeats and each sample within each biological replicate corresponds to a sample pooled from large populations in **a**, **d**, **e**; each biological replicate corresponds to a sample pooled from from 40-60 animals in **b**, **c**; every dot stands for averaged value from 3 technical replicates in **b**, **c** or strands for every technical replicates in **a**, **d** and **e**); Two-tailed Student’s t test was used in **b**, **c**; error bars mean ± S.E.M. *p<0.05, **p<0.01, ***p<0.001.

Unlike ATFS-1 and POLG, LONP-1 interacts equally with both wildtype mtDNAs and ΔmtDNAs (Fig. 5a and Supplementary Fig. 6c), suggesting heteroplasmy is not maintained by uneven mtDNA binding by the protease. We next examined the effect of inhibiting LONP-1 on heteroplasmy. Strikingly, *lonp-1* inhibition via RNAi increased wildtype mtDNAs (Fig. 5b) and reduced ΔmtDNAs, which improved the heteroplasmy ratio from 59% ΔmtDNAs to 25% (Fig. 5c). Similar results were obtained in *atfs-1^nuc(−)^* worms upon LONP-1 inhibition (Supplementary Fig. 6d). Importantly, *lonp-1*(RNAi) did not reduce the brood size of heteroplasmic worms (Supplementary Fig. 6e), suggesting that *lonp-1*(RNAi) did not select against embryos with dysfunctional mitochondria.

We next examined the interaction between ATFS-1 and mtDNAs in heteroplasmic worms upon *lonp-1* inhibition. Interestingly, when raised on *lonp-1*(RNAi), the percentage of ΔmtDNAs and wildtype mtDNAs bound to ATFS-1 nearly reflected the heteroplasmy percentage within the whole worm lysate (Fig. 5d). Thus, the mtDNA-bound protease is required to establish the enriched interaction between ATFS-1 and ΔmtDNAs. In addition to increasing the percentage of wildtype mtDNAs bound by ATFS-1, *lonp-1*(RNAi) also increased the percentage of POLG that interacted with wildtype mtDNAs (Fig. 5e, also see Fig. 4h). However, *lonp-1*(RNAi) did not alter the percentage of HMG-5/TFAM bound to ΔmtDNAs, consistent with HMG-5 interacting with all mtDNAs independent of replication (Supplementary Fig. 6f). In support of this interpretation, when heteroplasmic worms raised on *cco-1*(RNAi) or in *clk-1* mutant strain, wildtype mtDNAs were specifically increased (Supplementary Figs. 6g,h). Combined, these findings suggest that LONP-1-mediated proteolysis antagonizes the ability of mitochondrial ATFS-1 to stimulate mtDNA replication. LONP-1 activity may be compromised within mitochondrial compartments harboring ΔmtDNAs, leading to ATFS-1 accumulation and mtDNA replication. We propose that globally inhibiting LONP-1 promotes ATFS-1-mediated mtDNA replication throughout the cell, not just in compartments enriched in ΔmtDNAs, consequently leading to a recovery of wildtype mtDNAs and a depletion of ΔmtDNAs.

### LONP1 inhibition improves heteroplasmy and OXPHOS function in cybrid cells

Lastly, we examined if the role of LONP1 in maintaining ΔmtDNAs is conserved in mammals by examining two human heteroplasmic cybrid cell lines, which harbor a combination of wildtype mtDNA and ΔmtDNAs associated with mitochondrial disease^40^. While cybrid cells are often used as models of deleterious mtDNA heteroplasmy, it is important to note that cybrid cells are cancer cells in which patient-derived heteroplasmic mtDNAs were introduced by cell fusion. Thus, these lines may not replicate all aspects of deleterious heteroplasmy observed in affected patient tissues.

One cybrid line harbors a single nucleotide transition (COXI G6930A) that introduces a premature stop codon in the cytochrome c oxidase subunit I gene, which was isolated from a patient with a multisystem mitochondrial disorder (Fig. 6a)^41^. We also examined a cybrid line harboring a 4977 base pair deletion known as the “common deletion” which removes multiple OXPHOS genes and is associated with Kearns-Sayre Syndrome (KSS), progressive external ophthalmoplegia, cancer and aging (Fig. 6a)^42–45^. We first examined the impact of LONP1 siRNA on heteroplasmy in the KSS cybrid line. Similar to inhibition of *C. elegans* LONP-1, inhibition of human LONP1 by siRNA for 4 days (Fig. 6b), resulted in a 1.5-fold increase of wildtype mtDNAs (Fig. 6c) while KSS mtDNAs were decreased ~2 fold (Fig. 6d), resulting in a shift in the heteroplasmy ratio from 57.5% to 25.6% (Fig. 6d). This result suggests that the role of LONP1 in promoting heteroplasmy is conserved in mammals.

**Fig. 6.**
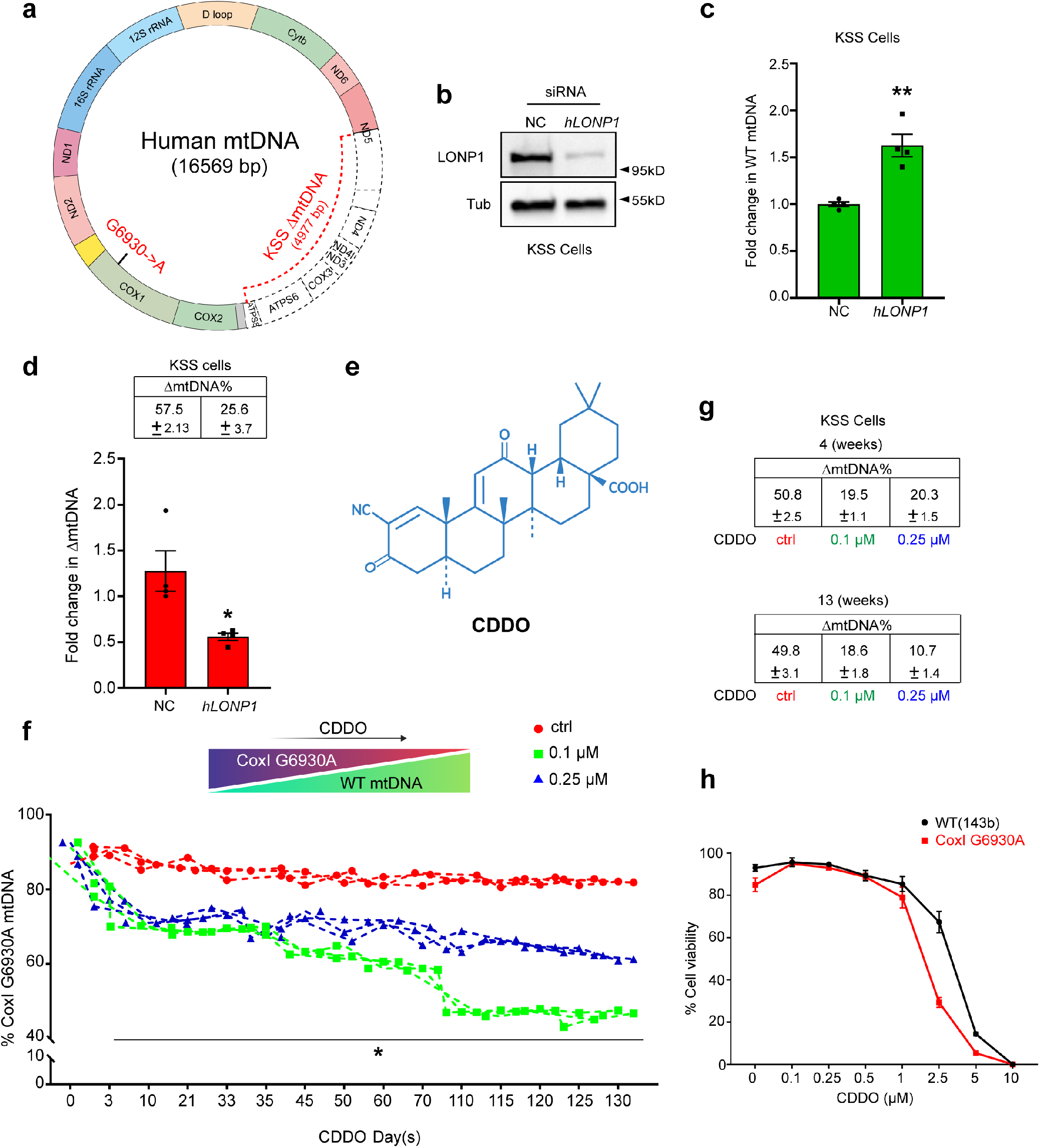
LONP1 inhibition improves heteroplasmy and OXPHOS function in cybrid cells. **a**, Schematic comparing human wildtype, KSS deletion (ΔmtDNA) and CoxI G6930A mtDNAs. **b**, LONP1 immunoblots from KSS heteroplasmic cells treated with *hLONP1* or control (NC) siRNA. Tubulin (Tub) serves as a loading control. **c**, WT mtDNA quantification in KSS cells treated with control or *hLONP1* siRNA (*n* = 3-4). **d**, Quantification of KSS ΔmtDNA in cells treated with control or *hLONP1* siRNA (*n* = 3). **e**, Chemical structure of CDDO. **f**, Quantification of G6930A mtDNA percentage following treatment with DMSO, 0.1 μM CDDO, or 0.25 μM CDDO at the indicated time points up to 130 days. *p<0.05 (*n* = 3). **g**, ΔmtDNA quantification in KSS heteroplasmic cells treated with DMSO, 0.1 μM CDDO, or 0.25 μM for 4 or 13 weeks. **h**, Cell viability of 143b(WT) and CoxI G6930A cells exposed to various concentrations of CDDO for 72 hours (*n* = 3 in 143b(WT) cell, *n* = 4 in CoxI G6930A cell). (“*n*” means independent biological replicates; every dot stands for averaged value from 3 technical replicates in **c**, **d**, **f** or stand for averaged value from biological replicates in **h**); Two-tailed Student’s t test was used in **c,d** and **f**; error bars mean ± S.E.M. *p<0.05, **p<0.01.

To further investigate the impact of LONP1 protease activity on heteroplasmy, we used the LONP1 inhibitor CDDO (2-cyano-3,12-dioxo-oleana-1,9(11)-dien-28-oic acid), also known as Bardoxolone (Fig. 6e)^20,46^. As determined by deep sequencing, the CoxI G6930A cybrid line initially harbored ~90% G6930A mutant mtDNAs and ~10% wildtype mtDNAs (Figs. 6a,f). Impressively, incubation with 0.1 μM or 0.25 μM CDDO for 3 weeks resulted in depletion of the CoxI G6930A mtDNA from 86% to 68% and 72%, respectively, with a concomitant increase in wildtype mtDNAs (Fig. 6f), Furthermore, continuous incubation with 0.1μM or 0.25 μM CDDO for 18.5 weeks further decreased the heteroplasmic ratios from ~90% to ~47% and 62%, respectively (Figs. 6f and Supplementary Fig. 7a). Similar results were obtained when the KSS cybrid cells were incubated with CDDO. As determined by qPCR, KSS cells initially harbored ~50% ΔmtDNAs. Incubation with 0.1 μM or 0.25 μM CDDO for 4 weeks and 13 weeks depleted ΔmtDNAs to 19.5%, 20.3 %, 18.6% and 10.7%, respectively (Fig. 6g). Importantly, neither 0.1 μM nor 0.25 μM CDDO affected viability or mitochondria respiration of the KSS, CoxI G6930A or 143b wildtype cell lines (Fig. 6h and Supplementary Figs. 7b,c) suggesting that the improved heteroplasmy ratio was not due to selection of cells with low levels of mutant mtDNAs.

Last, we determined the impact of the CDDO-dependent shifts in heteroplasmy on OXPHOS. Mitochondrial respiratory function was measured throughout the time course (Figs. 7a-d). Impressively, incubation of KSS cells with 0.1 μM or 0.25 μM CDDO for 4 or 13 weeks resulted in significant increases in basal respiration suggesting improved OXPHOS (Supplementary Fig. 7d). Furthermore, the improved heteroplasmy caused by CDDO in CoxI G6930A mtDNA also resulted in increased basal respiration and maximal respiratory capacity (Figs. 7b-d). For example, 3 weeks exposure to 0.1 μM CDDO improved basal oxygen consumption ~2-fold, while exposure for 18.5 weeks improved basal oxygen consumption over 3-fold (Figs. 7b,c). Taken together, these findings suggest that inhibition of LONP1 improves deleterious heteroplasmy and recovers mitochondrial respiration. Notably, these phenotypes are independent of mtDNA length, as maintenance of mutant mtDNAs with either a large deletion or a single base pair substitution require LONP1 function.

**Fig. 7.**
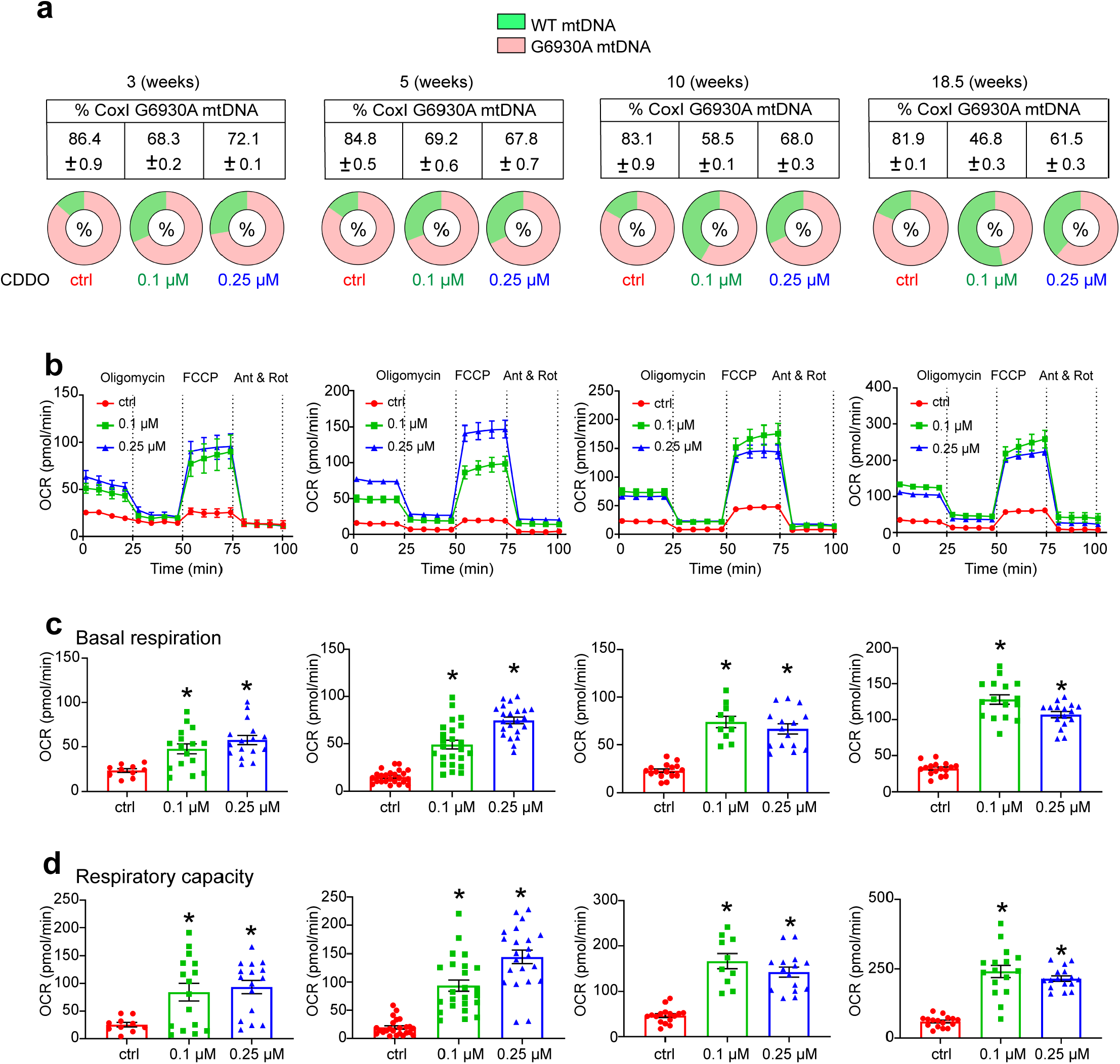
Pharmacological inhibition of LONP1 improves heteroplasmy and OXPHOS function in cybrid cells. **a**, Percentage of CoxI G6930A mtDNA in cells treated with DMSO, 0.1 μM or 0.25 μM CDDO for 3, 5, 10 or 18.5 weeks, *n* = 3; error bars mean ± S.E.M.; *p<0.05 (Student’s t-test). **b**, Oxygen consumption rates (OCR) of CoxI G6930A cells treated with DMSO, 0.1 μM or 0.25 μM CDDO for 3, 5, 10 or 18.5 weeks (*n* >=10). **c**, Quantification of basal respiration (*n* >=10). **d**, Quantification of respiratory capacity. (“*n*” means independent biological repeats; every dot stands for averaged value from 4 technical replicates in **c**,**d**); error bars mean ± S.E.M. *p<0.05.

## Discussion

The underlying mechanisms that govern heteroplasmy dynamics are largely unknown, however it has been proposed that mutant genomes have a selective advantage that promotes their accumulation. Previously, we and others reported that heteroplasmy depends on the transcription factor ATFS-1^9, 16^. In wildtype homoplasmic worms raised under normal conditions, ATFS-1 is imported into mitochondria and degraded by the protease LONP-1^17^. However, during mitochondrial dysfunction, a fraction of ATFS-1 accumulates within mitochondria and the nucleus. Moreover, we also reported that ATFS-1 binds to mtDNA within dysfunctional mitochondria^19^. Here, we report that mitochondrial-localized ATFS-1 promotes the accumulation of mtDNAs upon OXPHOS perturbation. Importantly, ATFS-1 and POLG binding to mtDNA was increased upon OXPHOS perturbation.

Deleterious heteroplasmy also causes OXPHOS dysfunction^47^ and increased mtDNA content which also required *atfs-1.* Furthermore, binding of ATFS-1 and POLG was also increased in heteroplasmic worms. Strikingly, ATFS-1 and POLG were both enriched on ΔmtDNAs by ~9:1 (Figs. 4g and 4h). As the ATFS-1 binding site is present in both wildtype and ΔmtDNAs, we propose that ATFS-1 accumulates within dysfunctional compartments, which in heteroplasmic worms harbor ΔmtDNAs. In support of this model, the nuclear activity of ATFS-1 is not required for the increased mtDNAs caused by *cco-1*(RNAi) in homoplasmic worms, or for maintaining ΔmtDNAs in heteroplasmic worms. Furthermore, nuclear ATFS-1 is not required for the increased POLG-mtDNA binding during OXPHOS perturbation in homoplasmic worms. Collectively, these results suggest that mitochondrial accumulation of ATFS-1 promotes recruitment of the mitochondrial replisome to mtDNA during OXPHOS dysfunction. It is important to note that additional regulatory mechanisms of mtDNA replication are likely involved during normal development as homoplasmic *atfs-1(null)* worms have only a slight reduction in mtDNAs (Fig. 3f).

We find that degradation of ATFS-1 by the mtDNA-bound protease LONP-1 is required to maintain heteroplasmy. In homoplasmic worms, knockdown of LONP-1 increases total mtDNA levels in an ATFS-1-dependent manner and increases POLG binding to mtDNA (Figs. 3f,g), consistent with a role for LONP-1 in regulating mtDNA replication^35, 48, 49^. In heteroplasmic worms, knockdown of LONP-1 also increases wildtype mtDNAs, but concomitantly reduces ΔmtDNA levels, improving heteroplasmy. Moreover, knockdown of LONP-1 abolishes the enriched binding of ATFS-1 and POLG to ΔmtDNAs (Figs. 5d,e). These results support a model wherein LONP-1 antagonizes the ability of ATFS-1 to promote mtDNA replication in healthy mitochondria. Because LONP-1 is an ATP-dependent protease that degrades proteins damaged by reactive oxygen species^31^, mitochondrial dysfunction may impede degradation of ATFS-1 by LONP-1, resulting in recruitment of POLG to mtDNA. We propose that this mechanism may serve to coordinate mtDNA replication with expansion of the mitochondrial network during cell growth or recovery from mitochondrial dysfunction. However, if compartmental dysfunction is caused by an enrichment of ΔmtDNAs, they are inadvertently, but preferentially, replicated. Our data suggests that inhibiting LONP1 throughout the mitochondrial network negates this preferential replication, leading to a reduction in the heteroplasmic ratio and recovery of mitochondrial function.

There are currently no FDA-approved treatments for diseases caused by mutant mtDNAs. However, the TORC1 inhibitor rapamycin has been shown to improve heteroplasmy in cybrid cells^50^. The findings indicate that inhibition of TORC1 in cybrid results in increased autophagy of dysfunctional mitochondria. It is perhaps interesting to note that inhibition of several TORC1 components also inhibit ATFS-1 function and *atfs-1*-dependent mitochondrial biogenesis in *C. elegans*^18^ suggesting TORC1 may function upstream of ATFS-1 in maintaining heteroplasmy. However, the role of TORC1 in regulating the function of mitochondrial-localized ATFS-1 remains to be determined. Here, we report that inhibition of LONP1 through siRNA-mediated knockdown or the small-molecule inhibitor CDDO reduces ΔmtDNA abundance in cybrid cells harboring patient-derived mutant mtDNAs with either a large deletion or a single base pair substitution. Moreover, this decrease in ΔmtDNA levels was accompanied by improved mitochondrial respiration in cybrid cells treated with CDDO. Our data suggests that LONP1 inhibition may represent a therapeutic strategy for diseases caused by mutant mtDNAs.

## ACKNOWLEDGMENTS

We thank the Caenorhabditis Genetics Center for providing *C. elegans* strains (funded by NIH Office of Research 362 Infrastructure Programs (P40 OD010440). We thank Carlos Moraes for the KSS and Giovanni Manfredi for the CoxI G6930A cybrid cell lines.

## Data and materials availability

All data in the paper are present in the paper or the Supplementary Materials. The ChIP-sequencing data have been deposited to the Gene Expression Omnibus database under the BioProject accession code PRJNA590136. The next-generation sequencing data have been deposited in the NCBI Sequence Read Archive database under the BioProject accession code PRJNA517630.

## AUTHOR CONTRIBUTIONS

Q.Y. and C.M.H. planned the experiments. Q.Y., Y.D., T.S., N.N., R.Z and P.C. generated the worm strains. Q.Y. performed the *C. elegans* and cybrid mtDNA analysis including ChIP and respiratory function. P.L., K.L. and S.W. performed and analyzed mtDNA sequencing. Q.Y., N.S.A. and C.M.H. wrote the manuscript.

## Supplemental Materials

Materials and Methods

Figures S1-S7

Tables S1-S2 (.xlsx)

References

