## Supplementary materials and figures for "mtDNA replication in dysfunctional mitochondria promotes deleterious heteroplasmy via the UPR^mt^"

**This PDF file includes:**

Materials and Methods
Figs. S1 to S7
References

**Other Supplementary Information for this manuscript include the following:**

Supplemental Table S1
Supplemental Table S2

### Materials and Methods

#### Worm strains

The reporter strain *hsp-6$_{pr}$::gfp* for visualizing $UPR^{mt}$ activation was previously described[1]. N2 (wildtype), and ΔmtDNA (or *uaDf5*) were obtained from the *Caenorhabditis* Genetics Center (Minneapolis, Minnesota). The *atfs-1(et18)* strain was a gift from Mark Pilon. The *atfs-1(null)*, or *atfs-1(cmh15),* strain was generated via CRISPR-Cas9 in wildtype worms as previously described[1]. The crRNAs (Integrated DNA Technologies) were co-injected with purified Cas9 protein, tracrRNA (Integrated DNA Technologies), and the *dpy-10* co-injection marker as described[2]. *atfs-1$^{nuc(-)}$* was introduced into both wildtype worms and the *hsp-6$_{pr}$::gfp* reporter strain via CRISPR-Cas9 (crRNAs and replacement sequence listed in Supplementary Table 1). *lonp-1*$^{FLAG}$ was introduced into both wildtype worms and the *hsp-6$_{pr}$::gfp* reporter strain via CRISPR-Cas9. Each strain was outcrossed at least 5 times. Unless otherwise noted, all worms were harvested between the late L3 and early L4 stages. All strains were maintained at 20°C.

#### *C. elegans* mtDNA and human cybrid cell KSS mtDNA quantification

L4 wildtype or *uaDf5* worms were placed on agar plates seeded with control(RNAi) or RNAi specific to the described OXPHOS genes and the F1 generation was harvested at the L4 stage. Wildtype mtDNA or ΔmtDNA quantification was performed using qPCR-based methods as described previously[3]. 40–60 worms were harvested in 35 μl of lysis buffer (50 mM KCl, 10 mM Tris-HCl (pH 8.3), 2.5 mM MgCl2, 0.45% NP-40, 0.45% Tween 20, 0.01% gelatin, with freshly added 200 μg/ml proteinase K) and frozen at −80°C for 20 min prior to lysis at 65°C for 80 min. Relative quantification was used for determining the fold changes in mtDNA between samples. 1 μl of lysate was used in each triplicate qPCR reaction. qPCR was performed using the iQ™ SYBR® Green Supermix and the Biorad qPCR CFX96™(Bio-Rad Laboratories). Primers that specifically amplify wildtype or ΔmtDNA are listed in Supplementary Table 1, as are primers that amplify both wildtype and ΔmtDNA (Total mtDNAs). Primers that amplify a non-coding region near the nuclear-encoded *ges-1* gene were used as an internal control for normalization (Supplementary Table 1).

For human patient fibroblast cell lines, wildtype and ΔKSS primers were used to detect wildtype mtDNA or ΔKSS mtDNA. Primers that amplify a sequence within the B2M (Human β2 myoglobin) gene were used as an internal control for normalization. Absolute quantification was also performed to determine the percentage or ratio of KSS ΔmtDNA relative to total mtDNA (KSS ΔmtDNA and wildtype mtDNA) as previously described[3]. Primers that specifically amplify wildtype or ΔmtDNA are listed in Supplementary Table 1. Standard curves for each qPCR primer set were generated using purified plasmids individually containing approximately 1 kb of the mtDNA fragments specific for each primer set.

**Chromatin immunoprecipitation (ChIP)**

ChIP assays for ATFS-1 and LONP-1$^{FLAG}$ were performed as previously described[4]. Synchronized worms were cultured in liquid and harvested at early L4 stage by sucrose flotation. The worms were lysed via Teflon homogenizer in cold PBS with protease inhibitors (Roche). Cross-linking of DNA and protein was performed by treating the worms with 1.85% formaldehyde with protease inhibitors for 15 min. Glycine was added to a final concentration of 125 mM and incubated for 5 min at room temperature to quench the formaldehyde. The pellets were resuspended twice in cold PBS with protease inhibitors. Samples were sonicated in a Bioruptor (Diagenode) for 15 min at 4°C on high intensity (30s on and 30s off). Samples were transferred to microfuge tubes and spun at 15,000*g for 15 min at 4°C. The supernatant was precleaned with pre-blocked ChIP-grade Pierce™ magnetic protein A/G beads (Thermo Scientific) and then incubated with Monoclonal ANTI-FLAG® M2 antibody (Sigma, F1804) or Mouse mAb IgG1 Isotype Control (Cell Signaling Technology, G3A1) rotating overnight at 4°C. The antibody-DNA complex was precipitated with protein A/G magnetic beads or protein A sepharose beads (Invitrogen). After washing, the crosslinks were reversed by incubation at 65°C overnight. The samples were then treated with RNaseA at 37°C for 1.5 hour followed by proteinase K at 55°C for 2 hours. Lastly, the immunoprecipitated and input DNA were purified with ChIP DNA Clean & Concentrator (Zymo Research, D5205) and used as templates for qPCR or next generation sequencing.

**mtDNA-immunoprecipitation (mtDNA-ChIP) and mtDNA quantification**

mtDNA-immunoprecipitation assays were performed similarly to the previously described ATFS-1 ChIP assay described[4], however the lysates were not sonicated so that wildtype and ∆mtDNA could be quantified by qPCR. Synchronized worms were cultured in liquid and harvested at early L4 stage by sucrose flotation. The worms were lysed via Teflon homogenizer in cold PBS with protease inhibitors (Roche). Cross-linking of DNA and protein was performed by treating the worms with 1.85% formaldehyde along with protease inhibitors for 20 min at room temperature. Glycine was added to a final concentration of 125 mM and incubated for 5 min at room temperature to quench the formaldehyde. The pellets were washed twice in cold PBS with protease inhibitor. Samples were transferred to microfuge tubes and spun at 15,000*g for 15 min at 4°C. The supernatant was precleaned with pre-blocked ChIP-grade Pierce™ magnetic protein A/G beads (Thermo Scientific) and then incubated with the described antibodies rotating overnight at 4°C. The antibody-mtDNA complex was precipitated with protein A/G magnetic beads (Thermo Scientific) (LONP-1$^{FLAG}$) or protein A sepharose beads (Invitrogen) (for ATFS-1, POLG, TFAM or LONP-1 antibodies. Sonicated salmon sperm DNA was used to block non-specific DNA binding on beads). After washing, the crosslinks were reversed by incubation at 65°C overnight. The samples were then treated with RNaseA at 37°C for 1.5 hour and then proteinase K at 55°C for 2 hours. Lastly, the samples were purified with ChIP DNA Clean & Concentrator (Zymo Research, D5205) and used as templates for qPCR. The results were normalized by input and either non-specific rabbit IgG or mouse IgG1 was used as a negative control. Primers that amplify wildtype, ∆mtDNA, or all mtDNAs (Total mtDNAs) are in Supplementary Table 1[3].

**ChIP-seq analysis**

The DNA fragments were sequenced using MiSeq at the University of Massachusetts Medical School Deep Sequencing Core. The quality of the raw sequencing data was first evaluated with fastqc (0.11.5). (https://www.bioinformatics.babraham.ac.uk/projects/fastqc/), and then mapped to the *C. elegans* genome (ce10 from UC Santa Cruz) by Burrows-Wheeler Aligner (BWA MEM, BWA version 0.7.15) algorithm with the standard default settings[5]. Duplicate reads were removed using picard tools v1.96 (https://broadinstitute.github.io/picard/). Peaks were determined using MACS version 2.1 [6] with the no-model parameter. Input was used as a control for peak-calling. Both narrow Peaks and broad Peaks were called, and the bigwig files were generated with the signal as fold enrichment by macs2 following the procedure at https://github.com/taoliu/MACS/wiki/Build-Signal-Track. The final set of peaks was determined if the difference in intensity values of control sample and input had a significance level of p-value < 0.01. IGV [7] was used to view the peaks and signals. To identify candidate LONP-1 interacting motifs, the regions that were highly enriched were used as input for MEME (http://meme.sdsc.edu). MEME was run using the parameters minw=8, maxw=25, in two modes (zoops & anr) and the significant motifs (E-value >= 1e-01). A background model is used by MEME to calculate the log likelihood ratio and statistical significance of the motif.

**Target site SNP frequency analysis in CoxI G6930A cells by deep sequencing**

Library construction for deep sequencing was modified from our previous report[7]. Following CDDO treatment, cells were harvested at different time points and genomic DNA extracted.

Briefly, regions flanking the CoxI G6930A site were PCR amplified using locus-specific primers bearing tails complementary to the Truseq adapters as described previously[8]. 50-100 ng input genomic DNA (mtDNA included) was PCR amplified with Phusion High Fidelity DNA Polymerase (New England Biolabs):(98°C, 15 s; 67°C 25 s; 72°C 18 s) × 30 cycles. 1 μl of each PCR reaction was amplified with barcoded primers to reconstitute the TruSeq adaptors using the Phusion High Fidelity DNA Polymerase (New England Biolabs): (98°C,15 s; 61°C, 25 s; 72°C, 18 s) × 9 cycles. Equal amounts of the products were pooled and gel purified. The purified library was deep sequenced using a paired-end 150bp Illumina MiSeq run.

MiSeq data analysis for editing at target sites or off-target sites was performed using a suite of Unix-based-software tools. First, the quality of the paired-end sequencing reads (R1 and R2 fastq files) was assessed using FastQC (http://www.bioinformatics.babraham.ac.uk/projects/fastqc/). Raw paired-end reads were combined using paired end read merger (PEAR)[9] to generate single merged high-quality full-length reads. Reads were then filtered by quality (using Filter FASTQC[8]) to remove those with a mean PHRED quality score under 30 and a minimum per base score under 24. Each group of reads was then aligned to a corresponding reference sequence using BWA (version 0.7.5) and SAMtools (version0.1.19). To determine background SNP or sequencing errors, all reads from each negative control replicate were combined and aligned, as described above. Background SNP types and frequencies were then cataloged in a text output format at each base using bam-readcount (https://github.com/genome/bam-readcount). For each drug treatment group, the average background SNP frequencies (based on SNP type, position and frequency) of the triplicate negative control group were subtracted to obtain the accurate SNP frequencies.

**QuantStudio 3D Digital PCR**

The detailed method was described previously[10]. All primers and probes were ordered from IDT (Coralville, Iowa). The 3D digital PCR was used according to the manufacturer's protocol. QuantStudio™ 3D Digital PCR Master Mix v2 and individual QuantStudio 3D digital PCR 20K Chip kit v2 were purchased from Thermo scientific (Applied Biosystems, Waltham, MA). Prepared sample mix was loaded into ChIP using QuantStudio 3D Digital PCR Chip Loader (Thermo scientific). Chip PCR amplification was performed in a ProFlex PCR System (96 °C for 10 min; 39

cycles of 60 °C for 2 min and 98 °C for 30 sec; and 60 °C for 2 min. After amplification, chips were loaded into QuantStudio 3D Digital reader to obtain results. Data analysis were analyzed using QuantStudio 3D Analysis Suite (Thermo scientific). All biological repeats were performed in at least triplicate to determine error.

∆mtDNA primers F: TTGCTTTTTCTTTATATGTTTTG; R: TTTATTTAATTTGGTTAAACAAGAGGT. ∆mtDNA probes 5' 6-FAM/ZEN/3' IBFQ: /56-FAM/AGGATCGTA/ZEN/ACATTTTATTTTTTTGCTTTA/3IABkFQ/.

Wildtype primers F: GCTTTTTCTTTATATGTTTTGTG.

R: TCACCTTCAGAAAAATCAAATGG wildtype mtDNA probes 5' HEX/ZEN/3' IBFQ: /5HEX/AATTATAGT/ZEN/AATTGCTGAACTTAACCGGGC/3IABkFQ/

**RNA isolation and qRT-PCR**

Total RNA was isolated from worm pellets using the TRIzol™ Reagent (Invitrogen). cDNA was then synthesized from total RNA using the iScript cDNA Synthesis Kit (Bio-Rad). qPCR was performed to determine the expression levels of the indicated genes using iQ™ SYBR GREEN supermix (Bio-Rad). Primer sequences are listed in Supplementary Table 1. Relative expression of target genes was normalized to the control. Fold changes in gene expression were calculated using the comparative CtΔΔCt method as previously described[4].

**Chemicals and Antibodies**

CDDO (Cayman Chemicals Cat No 81035). ATFS-1 polyclonal antibodies were generated and validated previously[11]. Polyclonal antibodies were generated to amino acid amino acids 1054-1072 of *C. elegans* POLG and subsequently affinity purified by Thermo Fisher Scientific Inc. Polyclonal antibodies were generated to amino acid amino acids 191-204 of *C. elegans* HMG-5 (TFAM) and subsequently affinity purified by Thermo Fisher Scientific Inc. Polyclonal antibodies were generated to amino acid amino acids 953-971of *C. elegans* LONP-1 and subsequently affinity purified by Thermo Fisher Scientific Inc. Monoclonal anti-FLAG® M2 antibody (Sigma, Cat # F1804), α-tubulin (Sigma), NDUFS3 (NUO-2 in *C. elegans*, complex I, Abcam). Supplementary Table 2.

**Cell Culture**

The KSS cell line was a gift from Carlos Moraes[12, 13]. The CoxI C6930A cell line was a gift from Giovanni Manfredi[14]. Cells were cultured in DMEM (4mM L-glutamine, 4.5 g/L glucose; Gibco, Thermo Fisher Scientific) plus 10% FBS with 1% pen-strep. Total cellular mtDNA was prepared as described[15]. Cells were incubated continuously in the described concentration of CDDO for the indicated number of days. The cells were sub-cultured prior to confluence every 48 hours.

**Cell Viability**

At the indicated time points, cells were stained with trypan blue[16] and quantified with an automated cell counter TC-20Tm (Bio-Rad). The results are an average of three independent assays.

**siRNA**

Cells were grown in 6-well plates and siRNAs were transfected with Lipofectamine RNAiMAX (Thermo Fisher Scientific Cat No 13778150) following the manufacturer's instructions. Human LONP1 siRNA was purchased from Dharmacon (L-003979-00-0005).

**Respiration Assays**

For mitochondrial respiration assays, oxygen consumption rate (OCR) was measured using a Seahorse Extracellular Flux Analyzer XFe96 (Seahorse Biosciences) as described[15]. 14,000 CoxI G6930A cells were seeded per well with fresh medium. OCR was measured using the Cell MitoStress Kit (as described by the manufacture). 180 µl of XF-Media was added to each well and then the plates were subjected to analysis following sequential introduction of 1.5 µM oligomycin, 1.0 μM FCCP and 0.5 μM rotenone/antimycin as indicated. Data is normalized to total protein as determined by the BCA protein assay.

**Western Blots and Mitochondrial Fractionation**

As previously described[17].

**Imaging and Fluorescence Quantification**

Whole worm images were obtained using either a Zeiss AxioCam MRm camera mounted on a Zeiss Imager Z2 microscope or a Zeiss M2BIO dissecting scope as described[1]. TMRE staining was performed by synchronizing and raising worms on plates previously soaked with S-Basal buffer containing DMSO, or final concentration 100 μM TMRE (Sigma, Cat No 87917). Prior to imaging, the TMRE-stained worms were transferred to plates seeded with control(RNAi) bacteria for 3 h to remove TMRE-containing bacteria from the digestive tract. Images were acquired using identical exposure times with a ZEISS LSM800 microscope with Airyscan. TMRE staining analysis is performed as described[18, 19]. In short, the average pixel intensity values were calculated by sampling images of different worms. The average pixel intensity for each animal was calculated using ImageJ (http://rsb.info.nih.gov/ij/). Mean values were compared using Student's t test or one-way (ANOVA) variance analysis followed by the post-hoc Tukey's test where appropriate. For each experiment, at least 20 worms were examined for each strain/condition. Statistical analysis was performed using the Prism software package (GraphPad Software).

**Statistical analysis**

All experiments were comprised of three independent biological replicates at least. Results were analyzed using Student's t test with a two-tailed distribution or one-way ANOVA (for multiple comparison) where appropriate, using GraphPad Prism software with corrected P values $< 0.05$ considered significant. All data are reported as mean ± SEM. Asterisks denote corresponding statistical significance *p < 0.05; **p < 0.01; ***p < 0.001 and ****p < 0.0001. The statistical tests performed and definition of ‘‘n’’ numbers in this study are indicated in the figure legends. For Figures 1d,1g-1h, 2c,2e-2i, 3b, 3e-3g, 4a,4e, 4g-4i, 5a-5e, 6c,6d, 6g-6h, 7a-7d, S1a, S2c, S2f-h, S3e, S4b, S4c, S4e-g, S5d, S5f, S6b-S6h, S7b-d, ‘‘n’’ means biological replicates. For C. elegans western blotting and gene expression, each sample within each biological replicate corresponds to a sample pooled from 3000-5000 animals. For cell culture western blotting, each sample within each biological replicate corresponds to one well from a tissue culture plate. For Figure 7b-d, “n” means biological replicates and each sample within each biological replicate corresponds to a sample pooled from 14000 cells in Figure 7b-d, 18000 cells in Supplementary Figure 7d (4 weeks) and 10000 cells (13 weeks). The deep sequencing to quantify heteroplasmy of the CoxI G6930A mtDNA following exposure to CDDO was double-blinded. Significance was accepted at p< 0.05. The researchers involved in experiments were not completely blinded during sample obtainment or data analysis.

Supplementary Figure 1

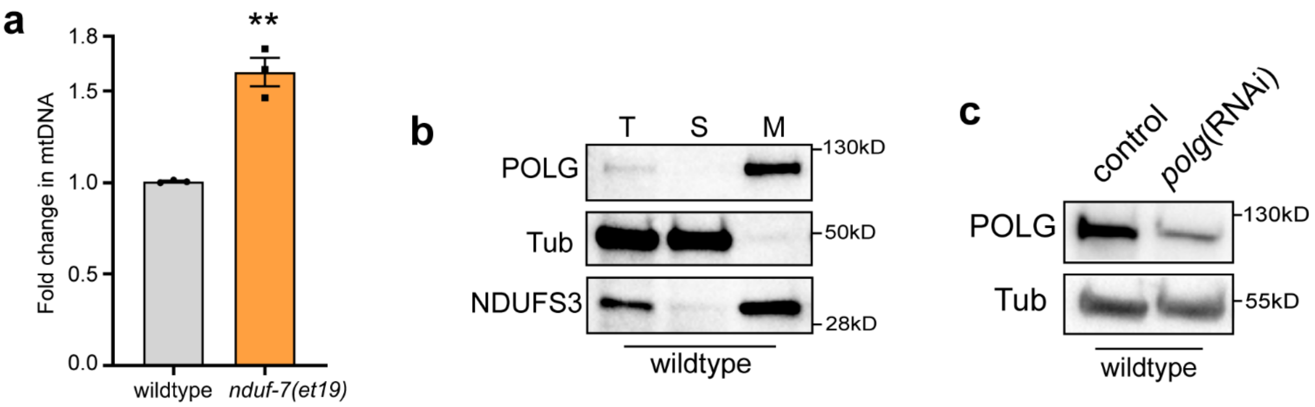


**Supplementary Fig.1 | Related to Fig. 1 and 2.**

**a**, Quantification of total mtDNA in wildtype and *nduf-7(et19)* worms (*n* = 3, Two-tailed Student's t test). **b**, POLG immunoblot of wildtype worms following fractionation into total lysate (T), post-mitochondrial supernatant (S), and mitochondrial pellet (M). Tubulin (Tub) and the OXPHOS protein (NDUFS3) serve as loading controls. **c**, POLG immunoblot of lysates from wildtype worms raised on control(RNAi) or *polg*(RNAi). Tubulin (Tub) serves as a loading control. ("*n*" means independent biological replicates and each sample contains 40-60 animals; every dot stands for averaged value from 3 technical replicates); error bars mean ± S.E.M. *p<0.05, **p<0.01.

Supplementary Figure 2

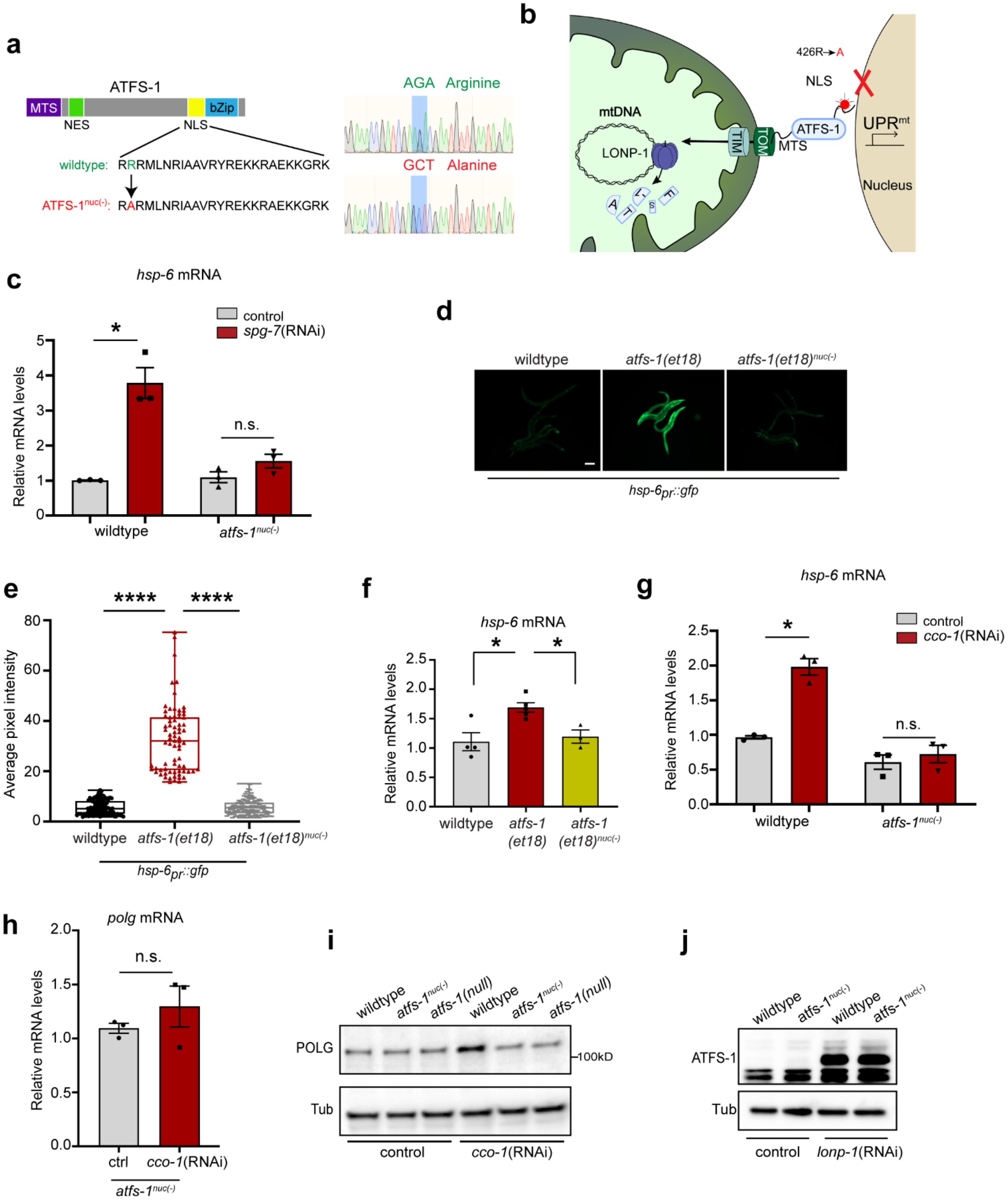

**Supplementary Fig. 2 | ATFS-1$^{nuc(-)}$ has an inhibited nuclear function. Related to Fig. 3.**

**a**, ATFS-1 schematic highlighting the R (Arginine) to A (Alanine) amino acid substitution to impair the nuclear localization sequence (NLS) within ATFS-1 yielding ATFS-1$^{nuc(-)}$. **b**, ATFS-1$^{nuc(-)}$/UPR$^{mt}$ signaling schematic with the arginine (R) to alanine (A) amino acid substitution at residue 426 that abolished the nuclear localization signal (NLS) in ATFS-1. **c**, Expression level of *hsp-6* mRNA in wildtype and *atfs-1$^{nuc(-)}$* worms raised on control(RNAi) or *spg-7*(RNAi) examined by qRT-PCR (*n* = 3, One-way ANOVA). **d-e**, Photomicrographs of wildtype, *atfs-1(et18)* and *atfs-1(et18)$^{nuc(-)}$;hsp-6$_{pr}$::gfp* worms (Scale bar 0.1 mm) (**d**); Quantification of fluorescence pixel intensity (Box & whiskers plots Min to Max) (**e**). **f**, Expression level of *hsp-6* mRNA in wildtype, *atfs-1(et18)* or *atfs-1(et18)$^{nuc(-)}$* worms examined by qRT-PCR (*n* = 3-5, One-way ANOVA). **g,** Expression level of *hsp-6* mRNA in wildtype and *atfs-1$^{nuc(-)}$* worms raised on control(RNAi) or *cco-1*(RNAi) examined by qRT-PCR (*n* = 3, One-way ANOVA). **h**, Expression level of *polg* mRNA in *atfs-1$^{nuc(-)}$* worms raised on control(RNAi) or *cco-1*(RNAi) examined by qRT-PCR (*n* = 3, Two-tailed Student's t test). **i**, POLG Immunoblots of lysates from wildtype, *atfs-1$^{nuc(-)}$* and *atfs-1(null)* worms raised on control or *cco-1*(RNAi). **j**, Immunoblots of lysates from wildtype and *atfs-1$^{nuc(-)}$* worms raised on control or *lonp-1*(RNAi). ATFS-1 or ATFS-1$^{nuc(-)}$ are indicated with an arrowhead. (In **c**,**f-h** "*n*" means independent biological replicates and each sample pooled from large populations; every dot stands for averaged value from 3 technical replicates); error bars mean ± S.E.M. *$p<0.05$, **$p<0.01$, ****$p<0.0001$.

Supplementary Figure 3

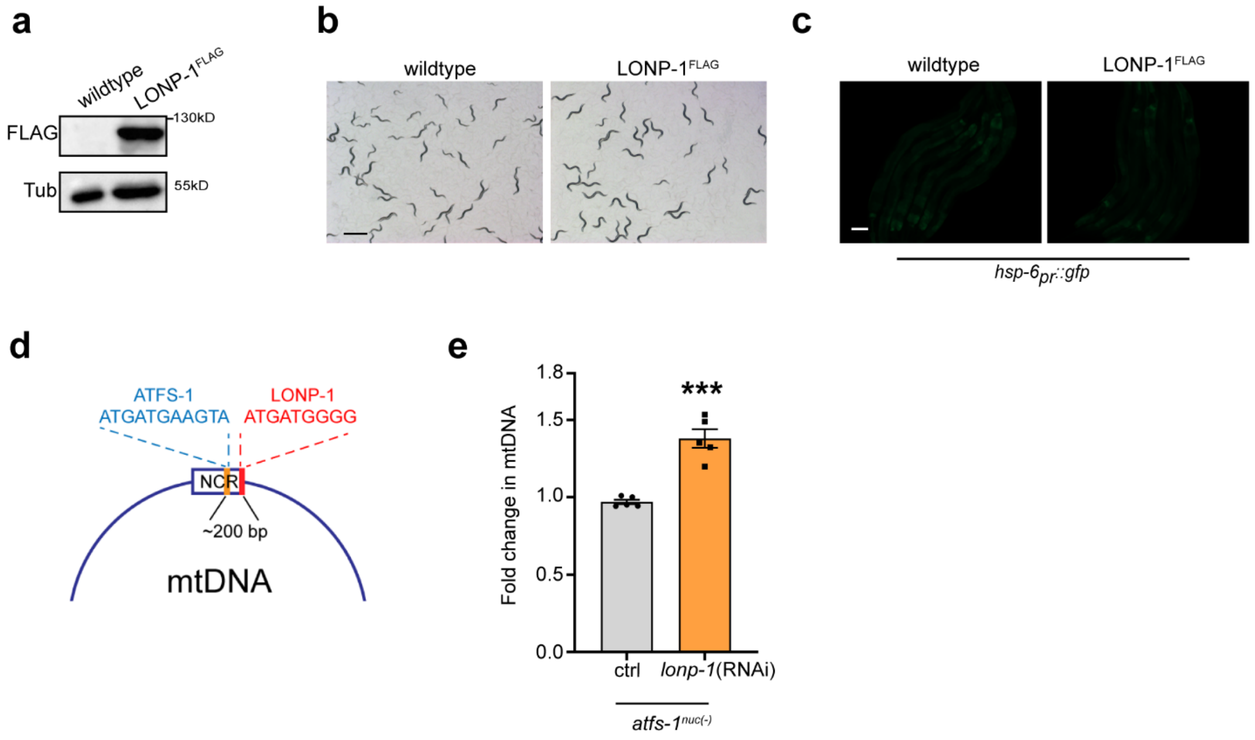


**Supplementary Fig. 3 | LONP-1$^{FLAG}$ does not affect worm development or UPR$^{mt}$ activation, Related to Fig. 3.**

**a**, FLAG immunoblots of lysates from wildtype and LONP-1$^{FLAG}$ wildtype worms. Tubulin (Tub) serves as a loading control. **b**, Images of wildtype or LONP-1$^{FLAG}$ worms 48 hours after synchronization indicating worms expressing LONP-1$^{FLAG}$ at the endogenous locus develop normally (Scale bar 1 mm). **c**, Fluorescent photomicrographs of wildtype *hsp-6*$_{pr}$*::gfp* or *lonp-1*$^{FLAG}$;*hsp-6*$_{pr}$*::gfp* worms 48 hours after synchronization indicating worms expressing LONP-1$^{FLAG}$ do not cause UPR$^{mt}$ activation. Scale bar 0.05 mm). **d**, Schematic of the putative ATFS-1 and LONP-1 binding sites within the mtDNA non-coding region (NCR) highlighting the proximity of both sites (~200 base pairs). **e**, Total mtDNA quantification in wildtype homoplasmic and *atfs-1*$^{nuc(-)}$ homoplasmic worms raised on control(RNAi) or *lonp-1*(RNAi) ($n$ = 5, Two-tailed Student's t test). ("$n$" means independent biological replicates and each sample contains 40-60 animals; every dot stands for averaged value from 3 technical replicates); error bars mean ± S.E.M. *p<0.05, **p<0.01, ****p<0.0001.

Supplementary Figure 4

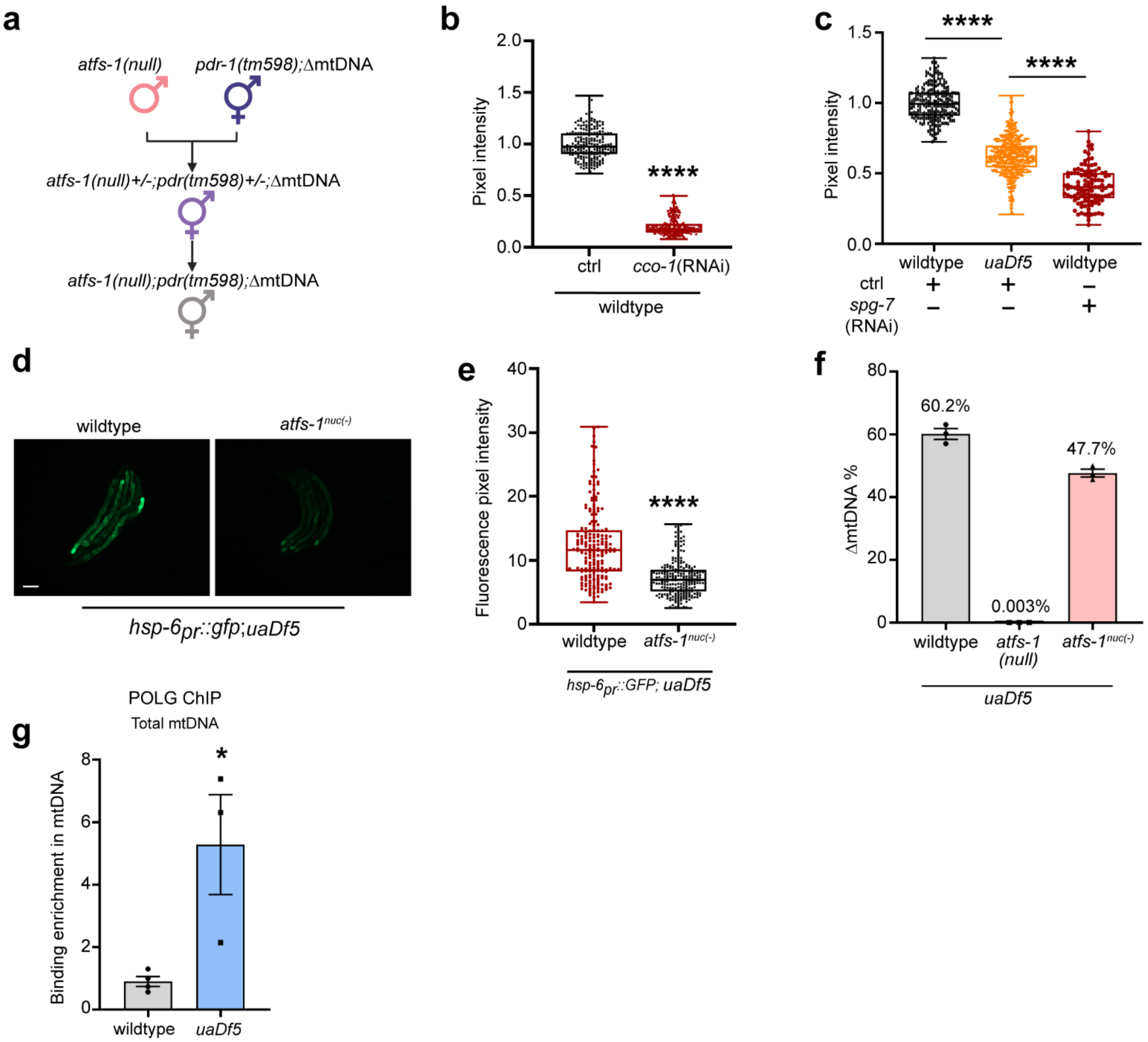

**Supplementary Fig. 4 | Unlike ATFS-1 and POLG, TFAM has no binding bias between ∆mtDNA and wildtype mtDNA, Related to Fig. 4.**

**a**, Crossing strategy of *atfs-1(null);pdr-1(tm598);uaDf5* strain. **b**, TMRE quantification of wildtype worms raised on control or *cco-1*(RNAi). **c**. TMRE quantification of heteroplasmic (∆mtDNA) worms raised on control(RNAi), or wildtype worms raised on control or *spg-7*(RNAi). **d,e**, Photomicrographs of *uaDf5* and *atfs-1*$^{nuc(-)}$*;uaDf5;hsp-6*$_{pr}$*::gfp* worms (Scale bar 0.1 mm) (**d**); Quantification of fluorescence pixel intensity (Box & whiskers plots Min to Max) (**e**). **f,** ∆mtDNA quantification as determined by qPCR in heteroplasmic *uaDf5* worms, *atfs-1(null)*;*uaDf5* worms and *atfs-1*$^{nuc(-)}$*;uaDf5* worms. **g**, Quantification of total mtDNA following POLG ChIP-mtDNA in homoplasmic wildtype or *uaDf5* worms (*n* = 3-4). (In **f**, "*n*" means biological replicates and each sample pooled from 40-60 animals; in **g**, "*n*" means independent biological replicates and each sample pooled from large populations; every dot stands for averaged value from 3 technical replicates in **f**,**g**); Two-tailed Student's t test was used in **b**,**e** and **g**; error bars mean ± S.E.M. *p<0.05, **p<0.01, ****p<0.0001.

Supplementary Figure 5

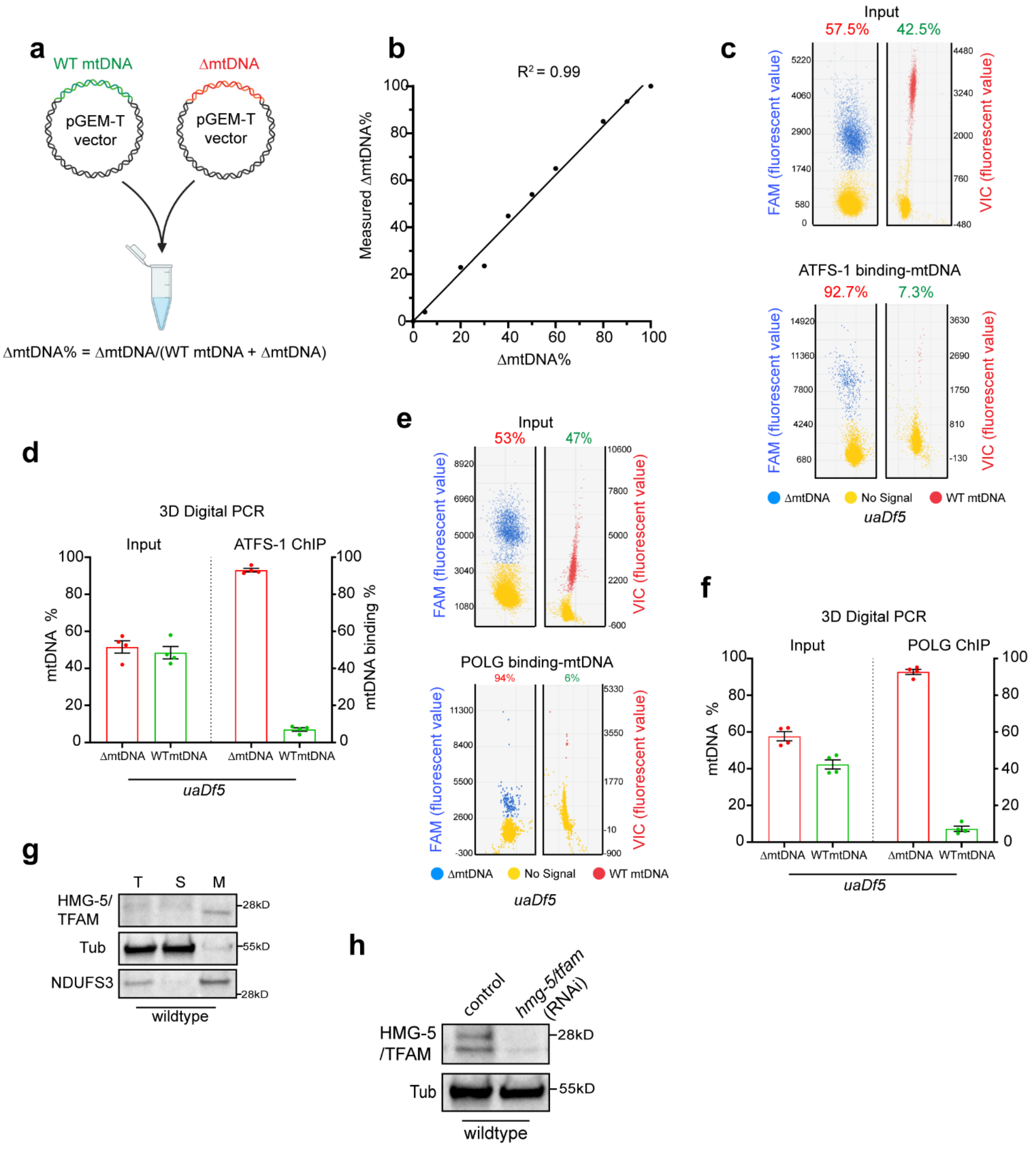

**Supplementary Fig. 5 | Related to Fig. 4**

**a**, Overview of qPCR strategy quantifying ΔmtDNA out of total mtDNA which is based on two specific pair of primers including ΔmtDNA and wildtype mtDNA primers[20]. **b**, Assay to determine the precision and accuracy of qPCR in quantifying the ratio of ΔmtDNA. They can amplify ΔmtDNA or wildtype mtDNA in same reaction respectively. **c-d**, Scatter plots (**c**) and results (**d**) of 3D digital PCR quantification of wildtype mtDNA and ΔmtDNA following ATFS-1 ChIP-mtDNA in heteroplasmic *uaDf5* worms (*n* = 4). **e-f**, Scatter plots (**e**) and results (**f**) of 3D digital PCR quantification of wildtype mtDNA and ΔmtDNA following POLG ChIP-mtDNA in heteroplasmic *uaDf5* worms (*n* = 4). **g**, HMG-5/TFAM immunoblot of wildtype worms following fractionation into total lysate (T), post-mitochondrial supernatant (S), and mitochondrial pellet (M). Tubulin (Tub) and the OXPHOS component (NDUFS3) serve as loading controls. **h**, HMG-5/TFAM immunoblots of lysates from wildtype worms raised on control or *hmg-5/tfam*(RNAi). Tubulin (Tub) serves as a loading control. ("*n*" means independent biological repeats and each sample pooled from large populations); error bars mean ± S.E.M. *p<0.05, **p<0.01, ****p<0.0001.

Supplementary Figure 6

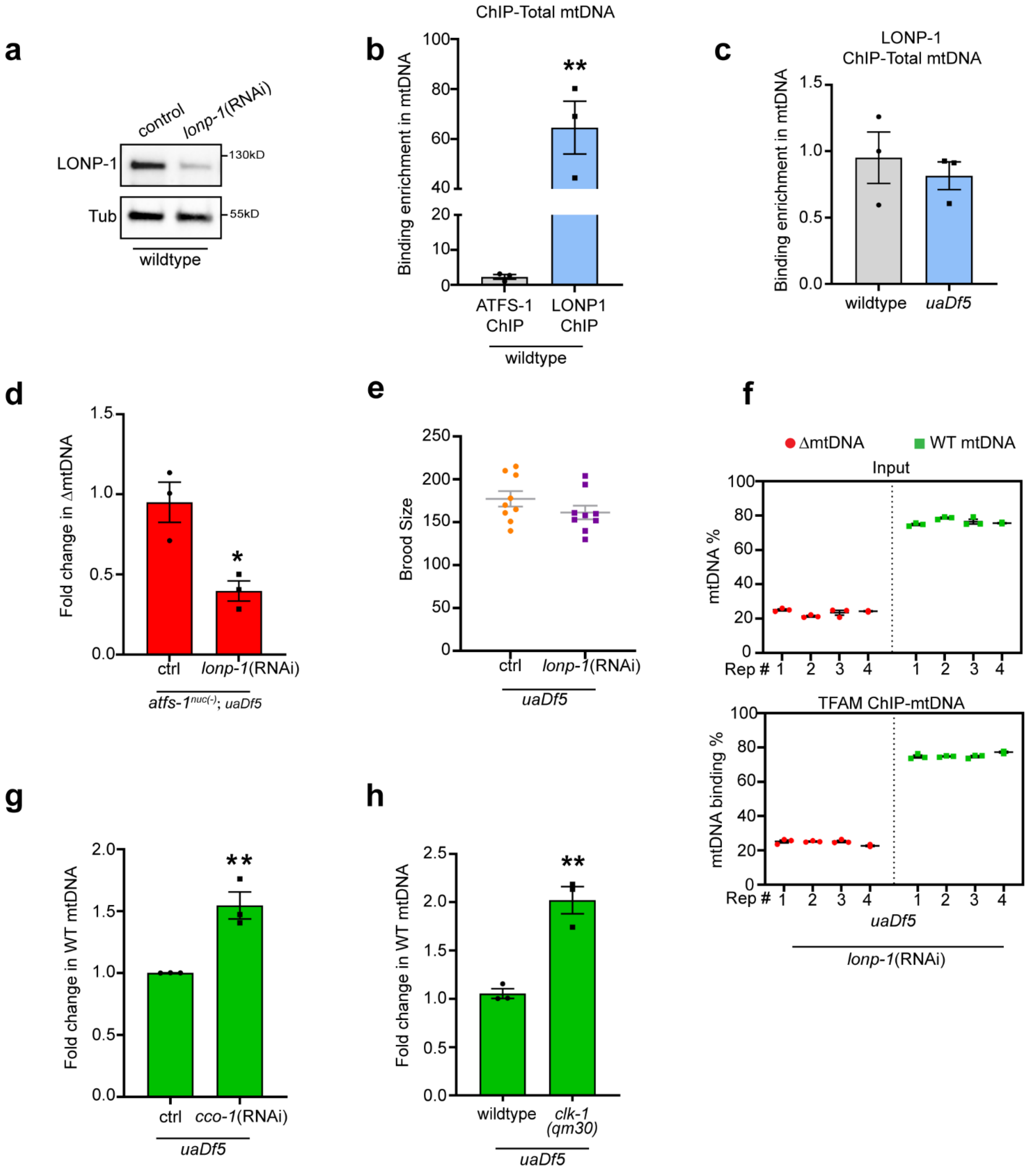

**Supplementary Fig. 6 | Inhibition of LONP-1 improves deleterious mtDNA level, Related to Fig. 5**

**a**, LONP-1 immunoblots of lysates from wildtype worms raised on control(RNAi) or *lonp-1*(RNAi). Tubulin (Tub) serves as a loading control. **b**, ChIP-mtDNA using ATFS-1 or LONP-1 antibodies in wildtype worms followed by quantification of total mtDNA ($n$ = 3). **c**, ChIP-mtDNA using LONP-1 antibodies in wildtype or heteroplasmic worms followed by quantification of total mtDNA (both wildtype and ΔmtDNA). **d**, ΔmtDNA quantification in *atfs-1*$^{nuc(-)}$*;uaDf5* worms raised on control(RNAi) or *lonp-1*(RNAi) ($n$ = 3). **e**, The brood size of heteroplasmic worms raised on control or *lonp-1*(RNAi) ($n$ = 9). **f**, ΔmtDNA and wildtype mtDNA quantification following HMG-5/TFAM ChIP-mtDNA in *uaDf5* heteroplasmic worms raised on *lonp-1*(RNAi) indicating that the binding HMG-5/TFAM to wildtype mtDNAs or ΔmtDNAs is the same as the input ($n$ = 4). **g**, wildtype mtDNA quantification in *uaDf5* heteroplasmic worms raised on control(RNAi) or *cco-1*(RNAi) ($n$ = 3). **h**, wildtype mtDNA quantification in *uaDf5* or *clk-1(qm30)*;*uaDf5* heteroplasmic worms ($n$ = 3). (In **b**,**c** and **f**, "*n*" means independent biological repeats and each sample pooled from large populations; in **d**,**g**,**h** "*n*" means independent biological replicate and each sample pooled from 40-60 animals; every dot stands for averaged value from 3 technical replicates in **b-d** and **g-h**); Two-tailed Student's t test was used; error bars mean ± S.E.M. *p<0.05, **p<0.01, ***p<0.001.

Supplementary Figure 7

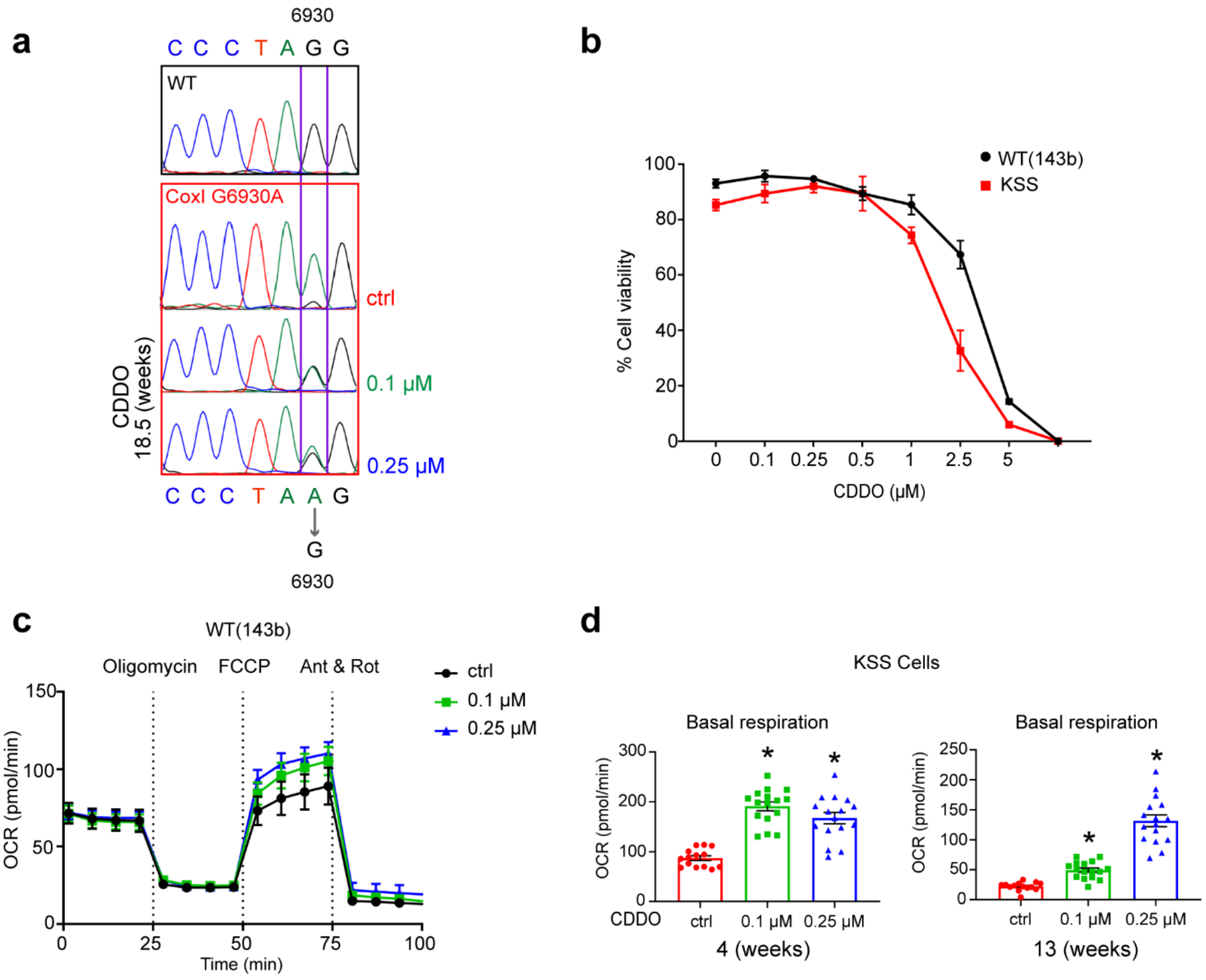


**Supplementary Fig. 7 | Pharmacological inhibition of LONP1 improves heteroplasmy and OXPHOS function in heteroplasmic cybrid cells, Related to Fig. 6 and 7.**

**a**, Confirmation of CDDO treatment by Sanger sequencing in CoxI G6930A cell. **b**, Cell viability of 143b(WT) and KSS ΔmtDNA cells exposed to various concentrations of CDDO for 72 hours (*n* = 3 in 143b(WT) cell, *n* = 4 in KSS cell). **c**, Oxygen consumption rates (OCR) of 143B (wildtype) cells treated with DMSO, 0.1 μM or 0.25 μM CDDO for 3 days (*n* = 22 or 24). **d**, Basal respiration of KSS heteroplasmic cells treated with DMSO (ctrl), 0.1 μM or 0.25 μM CDDO for 4 or 13 weeks (*n* = 14-16). ("*n*" means independent biological replicates; every dot stands for averaged value from biological replicates in **b** and **c**; Two-tailed Student's t test was used; error bars mean ± S.E.M. *p<0.05.

**Table S1.**

Primers and gRNAs for qPCR, qRT-PCR, mtDNA quantification and gene editing.

**Table S2.**

Supplemental Table 2. Antibodies

**Sequencing data**

All data in the manuscript are present in either the manuscript or the Supplementary Materials. The ChIP-sequencing data have been deposited to the Gene Expression Omnibus database under the BioProject accession code PRJNA590136. The next-generation sequencing data have been deposited in the NCBI Sequence Read Archive database under the BioProject accession code PRJNA517630.
